# Gelatinase activity is required for Müller glia morphogenesis and neuronal maintenance in the retina

**DOI:** 10.64898/2025.12.17.694830

**Authors:** Natalia Jaroszynska, Aanandita A. Kothurkar, Dylan Jankovic, Marybelle Cameron-Pack, Ryan B. MacDonald

## Abstract

Glial cells are indispensable support elements of the nervous system, yet the mechanisms by which they integrate into densely packed, extracellular matrix (ECM)-rich cellular environments remain poorly understood. In the retina, Müller glia (MG) serve as the principal support cells, performing roles analogous to astrocytes in the brain. As late-born retinal cells, MG undergo morphogenesis within a complex extracellular milieu composed of neighbouring neurons, synapses, and ECM components. Using the zebrafish retina, we show that developing MG dynamically interact with the ECM across all retinal layers while expressing the ECM-remodelling enzymes matrix metalloproteinases (Mmp) 2 and 9. Although Mmps are known to contribute to tissue remodelling and disease processes such as glioma invasion, their developmental roles in glia remain less defined. Here, we demonstrate that Mmp2 and Mmp9 are required for MG morphogenesis and glia–neuron integration. Pharmacological inhibition or genetic loss of *mmp2* and *mmp9* impaired MG outgrowth and reduced process complexity within synaptic layers, leading to decreased MG–synapse associations. Double mutants (*mmp2^−/−^;mmp9^−/−^*) displayed normal neuronal development and visual function at larval stages but exhibited gliosis, premature neurodegeneration, and visual deficits in adulthood. Together, these findings reveal that Mmp2 and Mmp9 are critical for establishing MG–synapse interactions during development and for maintaining neuronal integrity and visual function in the mature retina.

## Introduction

During development, neurons and glia must come together to make precise contacts and establish a partnership that is necessary for a correctly functioning central nervous system (CNS). Glial cells have elaborate morphologies that facilitate their ability to contact and integrate with their cellular partners, namely neurons and the vasculature^1^. A major point of glial-neuronal contact is at the synapse, where neuronal pre- and post-synaptic terminals are ensheathed by glial projections to form a functional compartment known as the tripartite synapse^2,3^. These contacts facilitate glia to provide essential support to their synaptic partners by expressing specific molecules necessary for energy metabolism, neurotransmitter recycling and ion homeostasis^4^. Altered glial morphology is a common pathological feature of many neurological disorders, as well as aging^5^, potentially disrupting their support functions and exacerbating neuronal degeneration. Despite their importance, little is known about how glial cells elaborate their complex morphology to form synaptic contacts during development, nor the consequences of disrupted glial contacts on neuronal maintenance in the mature CNS.

The retina, a component of the CNS, is composed of five major neuronal types: photoreceptors (PRs), amacrine cells (Acs), retinal ganglion cells (RGCs), bipolar cells (BPCs) and horizontal cells (HCs) and a single principal macroglia, the Mulller glia (MG)^6^. MG are considered molecular and functional homologues to astrocytes, performing analogous functions^7^. These retinal cells are organised into three discrete cellular layers, separated by two synaptic neuropils, forming relatively simple circuits required for vision^6^ (**Fig.1**). MG span all retinal layers and extend numerous morphological protrusions that are essential for their ability to contact neurons and perform their supportive functions. Most synapses in the retina are found in the inner plexiform layer (IPL) (**Fig.1**), where distinct neuronal types and MG form precise visual circuits segregated into discrete layers of synapses, or sub-laminae^8^. During development, nascent MG cells undergo dynamic morphogenesis to transform from simple unbranched radial cells to highly branched specialised cells to contact and support nearby neurons^8–10^. Similarly to other CNS tissues, the neurons in the developing retina are born first, subsequently followed by glia^10–14^. This means that developing MG face a dense extracellular environment consisting of other cells and the extracellular matrix (ECM) as they establish their elaborate morphology. We had previously conducted transcriptomics and a CRISPR-based screen which identified numerous molecules involved in MG morphogenesis, including members of the matrix metalloproteinase (Mmp) family, Mmp2 and 9^15^. However, their impact on MG morphogenesis and neuronal contacts has not yet been determined.

**Figure 1.**
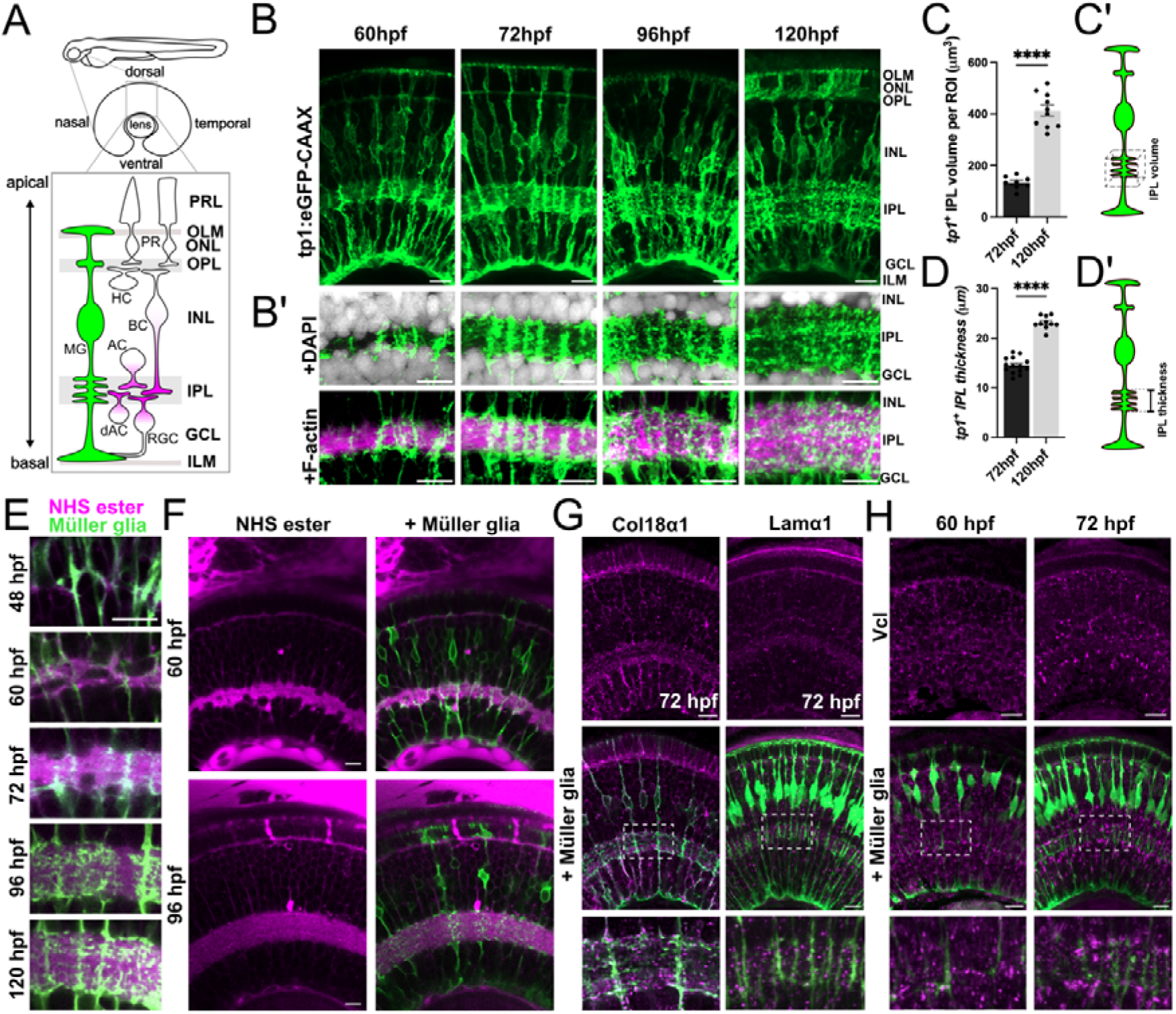
Müller glia develop in an ECM-rich extracellular environment. **(A)** Schematic diagram of the larval zebrafish retina indicating the dorsal region of the retina examined. Zoom box shows the cellular composition and organisation of retinal neurons and their processes (magenta) and Müller glia (MG; green). Photoreceptors, PR; horizontal cells, HC; bipolar cells, BC; amacrine cells, AC; displaced amacrine cells, dAC; retinal ganglion cells, RGC; Photoreceptor layer, PRL; outer limiting membrane, OLM; outer nuclear layer, ONL; outer plexiform layer, OPL; inner nuclear layer, INL; inner plexiform layer, IPL; ganglion cell layer, GCL; inner limiting membrane, ILM. **(B)** MG cell membranes labelled in the transgenic line Tg(*tp1:*eGFP-CAAX) between 60 and 120 hours post fertilisation (hpf). **(B’)** Close up of MG processes (green) in the IPL, surrounded by neuronal processes labelled with F-actin stain Phalloidin (magenta), nuclei labelled with DAPI (grey). **(C)** Measurement of MG process volume per region of interest (ROI) at 72 and 120 hpf, showing increasing complexity over time. **(C’)** Schematic indicating how volumetric measurements in (C) were performed. **(D)** Measurement of MG IPL process thickness at 72 and 120 hpf. **(D’)** Schematic of how measurements in D were performed. **(E)** NHS ester- AF646 (magenta) staining in the IPL surrounding MG processes (green) labelled by Tg(*tp1:*eGFP-CAAX). **(F)** Single z-plane images of NHS-ester staining and MG at 60 and 72 hpf. **(G)** IF with anti-Collagen18a (Col18a1) and anti-Laminin1 (Lama1) antibodies at 72 hpf; bottom panels show close up of the IPL. **(G)** Antibody staining for Vinculin (Vcl) at 60 and 72 hpf; bottom panels show close up of the IPL. Scale bars, 10µm.

MMPs are proteolytic enzymes that degrade both ECM and non-ECM substrates^16–20^. MMP-mediated ECM remodelling and cell–ECM interactions are crucial for shaping cell morphology in processes such as branching morphogenesis, angiogenesis, and astrocyte motility ^21^. In astrocytes, MMP2 activity regulates process complexity and responds to ECM stiffness ^22–24^. At the tips of cancer cell invadopodia, MMPs remodel the ECM to facilitate migration and invasion during epithelial–mesenchymal transition ^25–28^. MMP2 and MMP9, also known as the gelatinases, have been extensively linked to glioma progression, highlighting the interplay between these enzymes, the ECM, and glial cells ^29–31^. Dysregulated MMP activity and ECM alterations are also associated with age-related neurodegeneration and changes in mature glial morphology ^32–35^. In the retina, MMP2 and MMP9 are expressed in MG across species, both *in vitro* and *in vivo* ^36–40^. Loss of Mmp9 in zebrafish impairs MG-mediated PR regeneration ^41,42^, though its role in MG morphology under physiological conditions remains unclear. Studies on MMP2 have largely focused on vascular development ^43–45^, leaving the impact of the gelatinases on glial development poorly understood.

In this study, we investigated the consequences of the loss of gelatinase activity on glial development and maintenance. We focused on the morphologically complex MG cells in the retina and their interactions with retinal synapses in the major synaptic neuropil, the IPL. We assessed how MG navigate a dense environment to elaborate their fine processes and contact synapses during development. We identified that MG express active gelatinases during this morphogenesis period in the zebrafish retina. Pharmacological inhibition and genetic knockout of both gelatinases did not result in disrupted neuronal development or deficits in visual function in the embryonic retina. However, we observed a significant reduction in the morphological complexity of MG cells and glia-synapse contacts in the IPL in larval *mmp2^-/-^*;*mmp9^-/-^*double mutants. Impairing the activity or expression of a single gelatinase was insufficient to drive the same phenotypes in larval MG. Characterisation of gelatinase mutants in the adult retina revealed clear signs of vision loss and neurodegeneration across each retinal neuronal layer, including MG gliosis and inflammation. Thus, gelatinases are required for the proper establishment of glia-neuronal connections during development and neuronal maintenance in the mature CNS.

## Methods

### Animals

All adult zebrafish were raised in standard conditions under project license PP2133797. Zebrafish were maintained and bred at 28.5°C in a 14:10 light:dark cycle, and embryos were reared at 28.5°C until the desired developmental stage. Larvae were staged in hours post fertilisation (hpf), according to previously described methods^46,47^, and anaesthetized using 0.04% MS-222 (Sigma). To prevent pigmentation in whole-mount experiments, embryos were treated with 0.003% phenylthiourea (PTU; Sigma) for the duration of the experiment. Transgenic larvae used in this study are summarised in **Supplementary Table 1**.

### Immunofluorescence

Antibody staining was performed on zebrafish tissue fixed overnight in 4% paraformaldehyde (PFA). Following washes in PBS, fixed larvae or adult eyes were incubated in 30% sucrose overnight prior to cryosectioning. Antigen retrieval was performed on cryosections by heating the slides in sodium citrate (pH 6) for 20 minutes. Slides were subsequently washed in 0.1% PBS-Triton-X (PBSTx) and incubated in blocking solution (10% goat serum, 1% bovine serum albumin (BSA), in 0.1% PBSTx) for 1 hour at room temperature (RT). Slides were incubated with primary antibodies at 4°C overnight, followed by PBSTx washes, and incubation with secondary antibodies and DAPI stain for 2 hours at RT. Slides were washed again with PBSTx and mounted with glass coverslips overnight. Once dry, slides were imaged using a Zeiss confocal microscope with a 40x objective.

Whole-mount immunostaining was performed according to a previously published protocol^47^. Prior to staining, the larvae were treated with PTU to prevent pigment formation and aid imaging, as described above. Fixed larvae were washed in 1% PBSTween-20 (PBSTw), prior to antigen retrieval with Tris-HCl (pH 9) at 70°C for 15 mins. Larvae were washed extensively in PBSTw on a shaker and incubated in cold acetone for 20 minutes at -20°C. Following further washes, larvae were incubated in blocking solution (10% goat serum, 1% BSA, 1% Tween-20, in 0.1% PBSTx) for 2 hours at room temperature or at 4°C overnight. Larvae were then incubated in primary antibody solution for 2-5 days at 4°C on a gentle shaker, followed by extensive washes in 1% PBSTw. Next, larvae were stained with secondary antibody solutions for 2-3 days at 4°C on a gentle shaker. Once washed with PBSTw, larvae were embedded in 1% agarose in PBS on glass bottom dishes for confocal imaging. All antibody information and dilutions used are summarised in **Supplementary Table 2.**

### HCR

Hybridisation chain reaction (HCR) was performed using a protocol adapted from published protocols^48,49^. Probes against zebrafish *mmp14a* and *mmp14b* were designed using a custom python script from the Seuntjens lab (Github: https://github.com/SeuntjensLab/Easy_HCR) and obtained from ThermoFisher. All buffers used (hybridisation, wash, and amplification buffers) and HCR amplifiers (conjugated to Alexa Fluor^TM^) were obtained from Molecular Instruments (www.molecularinstruments.com). Freshly fixed tissues were used for HCR experiments to minimise degradation of mRNA in the tissue during long term storage. Larvae were then treated with 30 µg/mL Proteinase K for 10 mins on a shaker, followed by washes in 0.1% PBSTw and post fixation in 4% PFA for 20 mins at RT. After PBSTw washes, larvae were incubated in hybridisation buffer for 30 mins at 37 °C, prior to hybridisation of the probes at 37 °C overnight. Excess probe solution was removed by washing larvae with probe wash buffer at 37 °C, followed by washes with 5X SSCT at RT. Larvae were incubated in amplification buffer for 30 mins, and subsequently incubated with 12 pmol of amplifier hairpins, which were assembled by heating each hairpin at 95 °C for 90 seconds prior to this step. Excess hairpins were washed off with 5X SSCT at room temperature before confocal microscopy.

### NHS ester microinjection

Alexa Fluor™ 647 NHS Ester (Succinimidyl Ester) (Thermo Fisher Scientific; A20006) was used to label ECM proteins using the protocol adapted from Fischer et al^50^. The stock was diluted in 1X PBS and adjusted to pH 9. Fish at each stage were anesthetised in 0.04% MS-222 before being embedded in 1% low-melting point agarose. Fish were injected in the heart with 1 nL of 1X PBS or NHS Ester (pH 9) and incubated at 28°C for 30 minutes before confocal imaging.

### Fluorescent hybridising peptide (fCHP) staining

Cy3-conjugated fCHP (3Helix) staining for the detection of total and degraded fibrillar collagens was performed in whole-mount larvae according to published protocols^51,52^. fCHP was activated prior to use by heat shocking the 1X solution at 70°C for 5 minutes and quenching on ice immediately after. Larvae were subsequently incubated in the activated fCHP solution overnight at 4°C on a shaker and protected from the light. Negative control samples were incubated with non-activated fCHP, while brief heat shock of larvae prior to staining allowed for breakdown of fibrillar collagens and global fCHP incorporation to visualise total collagen. The solution was washed out extensively with PBS prior to confocal imaging.

### Pharmacological treatments

Larvae were treated with ARP-100 (Generon), AG-L-66085 (Sigma Aldrich), SB-3CT (Merck) or 0.1% DMSO (Thermo Fisher) at the 60 or 72 hpf by supplementing the E3 embryo media with the compounds as well as PTU. The concentrations of inhibitors used are listed in **Supplementary Table 3**. Where required, drugs were washed off after the treatment and transferred into fresh E3 media with PTU in a clean petri dish or multi-well plate prior to fixation.

### Gelatinase mutant generation

The approach used to generate stable knock out mutants using CRISPR-Cas9 mutagenesis was adapted from El-Brolosy et al.^53^. Guide RNAs (gRNAs) were designed using CHOPCHOP and acquired from Integrated DNA Technologies (IDT). 1-cell stage embryos were injected with Cas9 protein and *mmp2* or *mmp9* gRNAs, designed and prepared according to the protocol by Kröll et al.^54^. gRNA and genotyping primer sequences are listed in **Supplementary Table 4.**

### Optokinetic response (OKR)

For the OKR assay on 120 hpf-old larval zebrafish, the anaesthetised larvae were immobilised in 3% low melting point agarose prior to the measurements. Once set, the agarose was cut using a scalpel to allow for eye movement during the experiment and larvae were kept in E3 embryo media and left recover for 2 hours at 28.5 °C. Using a custom rig, a cylindrical screen with sinusoidal gratings of varying spatial frequencies were presented to the larvae, whose eye movements were tracked using a custom software. Eye velocity and other measurements were done using an adapted custom MATLAB script (original script available at https://bitbucket.org/biancolab/okrsuite), following stimulation with different spatial frequencies and contrast. For the adult OKR assay, zebrafish were anaesthetised and placed inside a custom- made sponge submerged in fish water. Fish were individually placed in the centre of the rotation chamber with 0.8 mm black and white stripes. The chamber was rotated at 8 rpm and the number of eye saccades was counted manually.

### Image analysis

All analysis was performed on 3D z-stacks using Fiji and/or Imaris software. For retinal layer thickness measurements and cell counts, Fiji line and multi-point tools were used, respectively. Regions of interest (ROIs) of 100µm x 100µm x 10 µm (xyz) were analysed, while for MG numbers, cells were counted in the whole field of view (160x 160x10µm ROI) due to their larger size and lower overall numbers. Retinal thickness measurements were performed by measuring the length of the apicobasal retinal axis based on DAPI staining in either cryosections or whole-mount retinas. Individual layer thickness was measured in the same way and normalised to the overall retinal thickness to obtain a ratio of each layer thickness compared to the retinal thickness, termed ‘Relative thickness’. The same approach was used for the measurements of MG IPL thickness, which was measured in Tg(*tp1*:eGFP-CAAX)-labelled retinas. All volumetric measurements were performed using Imaris by generating a surface to segment the object of interest and computing the volume within a ROI. For Ribeye-MG contact quantification, the ‘Spots’ module on IMARIS was used to identify individual puncta based on absolute intensity and compute their distance to MG process surfaces. ‘Close’ ribeye puncta were classified based on a distance threshold which was set as half of the mean diameter of Ribeye puncta and the percentage of ‘close’ spots from the total number of puncta in the ROI was subsequently quantified and compared between conditions. Gfap apicobasal distribution was quantified using Imaris, by generating a surface based on anti-Zrf1 (Gfap) IF staining and measuring the volume of the Gfap surface in the apical versus basal half of the cell. The ROIs for volumetric measurements were drawn based on the height of the MG cell from basal end-feet to the apical end of the cell, as labelled by anti-Zrf1. This ROI was then subdivided into 2 equal halves and volumes determined therein.

### RT-qPCR

Total RNA extraction from fresh-frozen whole larvae was performed using the RNeasy Mini kit (Qiagen) following manufacturer’s instructions and in RNAse-free conditions. For isolation of RNA from retinas at 72 hpf, larvae were first fixed with 4% PFA for 30 mins at room temperature followed by RNA extraction using the FFPE RNeasy Mini kit (Qiagen). cDNA synthesis was performed by reverse transcription of 1 µg mRNA using qScriptTM cDNA SuperMix (VWR) according to manufacturer’s protocol. 1:10 diluted cDNA was used as a template for Validated zebrafish-specific primers spanning exon-exon junctions used are listed in **Table S5.** Following RT-qPCR, raw CT values were normalised to the housekeeping gene actin (*actb1*) and relative expression was calculated as 2^-(MeanΔΔCt±SEMΔΔCt)^; Statistical analysis was performed using the ΔCt values.

## Results

### Müller glia undergo morphogenesis in a complex environment with a diverse ECM landscape

MG are amongst the last-born cells during retinal development^11^, and hence their morphological maturation takes place within a dense cellular environment^55^. A key morphological feature of MG is the numerous filopodial processes that extend into the neuropil layers, where they contact neuronal synapses to support neurotransmission **(Fig. 1A)**. To gain a better understanding of the full complexity of MG processes and their interactions with the local environment, we used the transgenic line Tg(*tp1:*eGFP-CAAX)*^u911^* to visualise the membranes of fine processes within the IPL throughout MG morphological maturation^56^ (**Fig. 1B-B’**). Co-labelling with phalloidin, a label for F-actin, allowed us to simultaneously visualise MG processes and those of the surrounding neurons (**Fig.1B**). As previously reported^9,56^, we observed a progressive increase in the complexity of MG processes between 60 and 120 hpf, with the majority of elaboration occurring between 60-96hpf, as evidenced by their increasing volume and the overall thickness of this layer over time (**Fig.1C-D’**). This expansion of MG processes and overall IPL thickness was also accompanied by an increase in phalloidin staining in this layer (**Fig.1B’**), indicative of the simultaneous increase in actin-rich neuronal and glial projection complexity.

The ECM has been implicated in retinal development and synaptogenesis ^57^. However, the direct interactions of MG cells with the ECM and its influence on their development have not been investigated. To assess the overall ECM landscape during MG morphogenesis, we used a fluorescently conjugated N-hydroxysuccinamide (NHS) ester, which globally labels ECM proteins in the extracellular space by binding their amine groups^50,58^. Following NHS ester injection, we evaluated its localisation in the retinas of live Tg(*tp1:*eGFP-CAAX) larvae (**Fig.1E,F; Fig.S1A**). NHS ester labelling could be seen in the extracellular spaces throughout the retina and surrounding extraretinal tissues (**Fig.1F**;**Fig.S1A)**. The most prominent signal was detected within the synaptic layers, particularly the IPL, into which MG dynamically extend their processes throughout morphogenesis (**Fig.1E**). We further assessed the retinal expression of all fibrillar collagens stained with fluorescent Collagen hybridising peptide (fCHP) but did not observe any notable expression or turnover of total collagen in the IPL, only the retinal basement membranes (ILM/OLM) (**Fig.S1B**).

To assess the ECM composition and MG interaction in more detail, we next performed immunofluorescence (IF) staining for several classical ECM components with spatial relationship with developing MG (**Fig.1G**;**Fig.S1C-E**). We detected several ECM markers in the IPL with varying temporal dynamics, suggesting that the ECM is still being laid down and remodelled during this time. For example, Collagen 18a1 immunoreactivity was observed in MG and both synaptic layers at 72 hpf, and from 96 hpf, it was exclusively detected in the OPL (**Fig.1G**;**Fig.S1C**). We also saw Laminin-alpha 1 localisation shift from only basement membrane to more diffuse expression across the neuroretina, including the IPL (**Fig.S1D**). Decorin, another ECM component, was found to mirror the OPL-specific expression of Col18a1 (**Fig.S1E**), suggesting that different ECM components may play distinct roles in discrete retinal layers. In addition to ECM components, we also observed the expression of focal adhesion protein Vinculin at the tips of nascent MG processes at 60 and 72 hpf (**Fig.1H**), indicative of direct MG-ECM interactions during this time.

Together, these observations demonstrate that developing MG cells encounter a range of different classes of ECM components in their local environment, which they must interact with and navigate to establish their mature morphology and integrate into neuronal circuits during development.

### ECM-remodelling enzymes Mmp2 and Mmp9 are expressed and activated in Müller glia undergoing morphogenesis

Having shown that MG cells develop in a dense extracellular environment, we next sought to understand how MG navigate the ECM during their dynamic morphological elaboration. Several studies have shown that mature MG produce the ECM-remodelling enzymes Mmp2/9, however their expression during development is not well described. To characterise their expression during MG morphogenesis, we performed IF with antibodies against Mmp2 and Mmp9 in the MG-specific transgenic reporter Tg(*tp1:*eGFP-CAAX) at developmental stages when MG are dynamically elaborating their morphology and interacting with their environment (**Fig. 2A,B** and **Fig.S2A,B**). While we observed diffuse staining throughout the retina for both Mmp2 and Mmp9 at both early and more advanced stages of MG morphogenesis, their expression was highly enriched in MG cells (**Fig.2A,B**). This was particularly apparent at MG outer membranes, as seen in the co-localisation with eGFP-CAAX (**Fig.2A, B** and **Fig.S2A,B)**. At 60 hpf, the onset of MG morphogenesis, we found that both gelatinases were present in the translocating cell bodies as well as around the nascent plexiform layer processes and MG end-feet, where they associate with the basement membrane (**Fig. 2A,B**). At later stages of development, Mmp2 and 9 immunoreactivity was still observed in the developing plexiform layers, particularly the IPL, correlating with the rapid extension of MG lateral processes into this layer (**Fig.2A,B** and **Fig.S2A,B;** dotted lines). Thus, both gelatinases are expressed by MG throughout their dynamic morphological development, including within the ECM-rich IPL.

**Figure 2.**
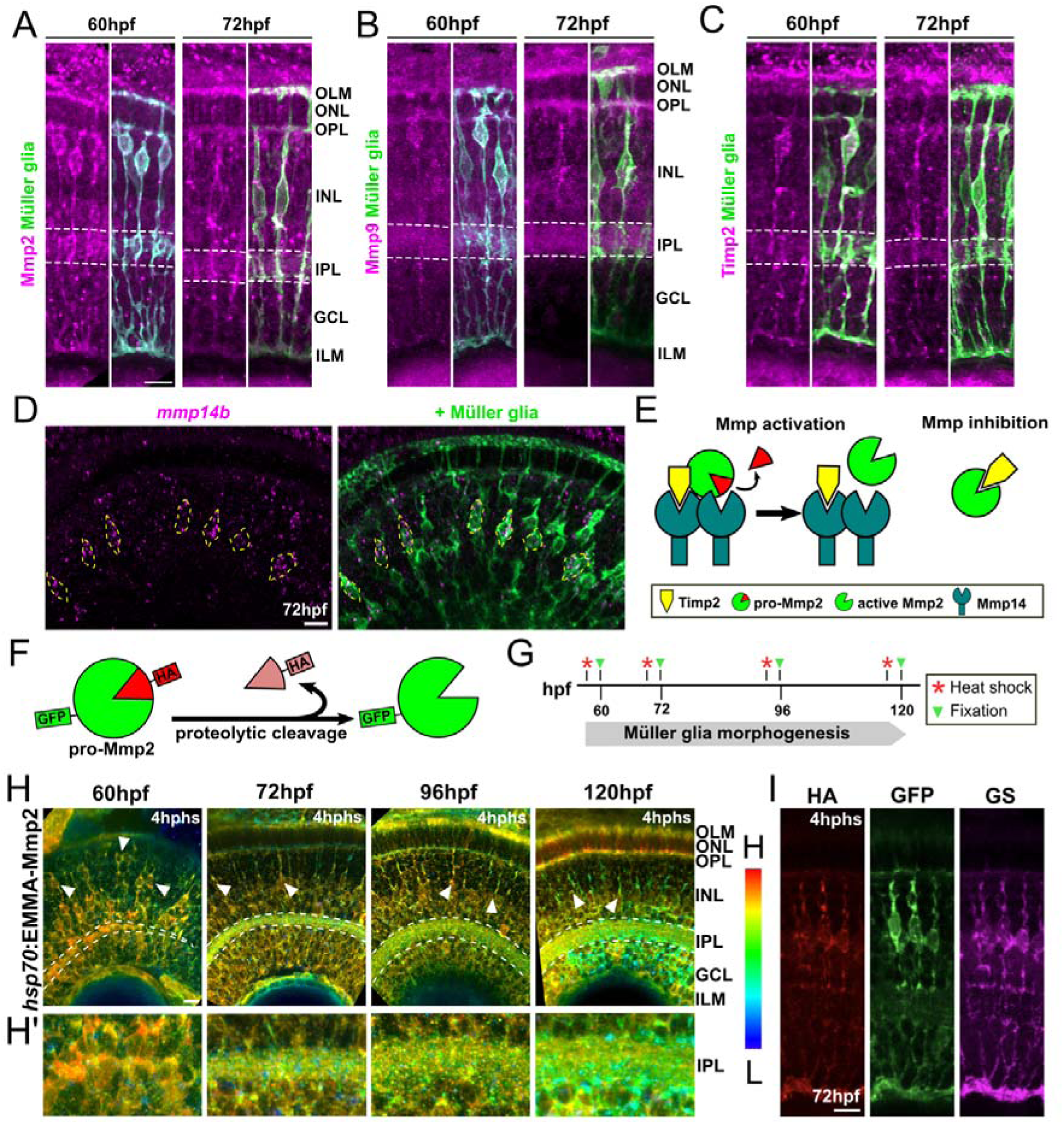
Gelatinases Mmp2 and Mmp9 are expressed and activated during Müller glial morphogenesis. **(A)** Whole mount immunofluorescence (IF) of Mmp2 (magenta) in Tg(*tp1:*eGFP-CAAX)-labelled MG (green) in the zebrafish retina at 60 and 72 hpf. Dotted lines indicate the IPL. **(B)** IF for Mmp9 (magenta) in MG cells (green) at 60 and 72 hpf. Dotted lines indicate the IPL. **(C)** IF for Timp2 (magenta) in MG cells (green) at 60 and 72hpf. **(D)** HCR *in situ* hybridisation detecting *mmp14b* mRNA in MG (green), labelled by Tg(*tp1:*eGFP-CAAX) at 72 hpf; yellow dotted lines indicate MG nuclei. **(E)** Schematic of MMP2 activity regulation. Pro-Mmp2 is activated by Mmp14 in complex with Timp2, through the proteolytic cleavage of the Mmp2 pro-domain. Active Mmp2 remains at the membrane bound to Mmp14 and is inhibited by Timp2. **(F)** Schematic of the Epitope Mediated MMP Activation (EMMA) reporter assay, where double-labelled Mmp2 is expressed under the control of heat shock-inducible promoter. **(G)** Summary of experiment: Tg(*hsp70:*EMMA-Mmp2) embryos heat shocked at 60, 72, 96 and 120 hpf, corresponding to different time-points during MG morphogenesis, and fixed 4 hours post heat shock (hphs) prior to IF for GFP and HA tag, as shown in (H). **(G)** EMMA-Mmp2 ratiometric images of HA/GFP, or activation maps, showing proportion of active (green-red) cleaved and inactive (blue) Mmp2 **(H’)** Close up of IPL region, showing Mmp2 activation throughout MG development. **(I)** IF for inactive (HA) and active (GFP) portions of Mmp2 and GS, a MG marker. Scale bars, 10µm.

Gelatinases require proteolytic cleavage to be activated, and therefore expression alone does not indicate Mmp2/9 activity^59^. While evidence in the chicken retina showed gelatinase activity in the retinal neuropil layers^40^, their activity has not been investigated during retinal development. First, to assess the capacity of MG to activate gelatinases, we performed IF and *in situ* hybridisation chain reaction (HCR) to assess the expression of key gelatinase activity regulators, tissue inhibitors of metalloproteinases 2 (Timp2), and *mmp14a* and *mmp14b* (**Fig.2C,D**). When in complex with Timp2, Mmp14 activates pro-Mmp2 by cleavage of the pro-domain, while Timp2 alone acts on active Mmp2 to inhibit it^60^. Antibody staining for Timp2, revealed its presence at the MG membranes simultaneously with gelatinase expression and activation (**Fig.2C;Fig.S2C**). We also detected *mmp14b* mRNA, but not *mmp14a* in the MG cell body during their morphogenesis, suggesting that the activating machinery for Mmp2 is also present at the time of MG morphogenesis (**Fig. 2D;Fig.S2D-E**). Next, to assess gelatinase activation patterns in the developing retina, we used the Epitope mediated MMP activation (EMMA) assay to visualise both the zymogenic pro-Mmp2 and its active, cleaved form **(Fig. 2E)** ^61^. Using the reporter line, Tg(*hsp70*:EMMAed-Mmp2), we performed heat-shock to drive the expression of EMMA-Mmp2 at different stages of MG development and assessed its activation pattern **(Fig. 2F).** At 60hpf, we observed an accumulation and activation of EMMA-Mmp2 in the apically located MG soma (**Fig. 2G, arrowheads**). Additionally, EMMA-Mmp2 activation was also detected in the presumptive IPL, surrounding developing RGCs and ACs and their projections (**Fig. 2G’**). At 72hpf, we observed reduced activation of EMMA-Mmp2 in the cell bodies, and increased activation within the plexiform layers as development proceeds. The identity of MG cells was confirmed by antibody staining for glutamine synthetase (GS), a specific marker for MG cells (**Fig.2H)**.

Together, these findings indicate that gelatinases are expressed and activated in and around developing MG cells, including their elaborating IPL processes.

### Functional redundancy and compensation between gelatinases mask phenotypes in loss of function *mmp2* and *mmp9* mutants

We next sought to examine whether gelatinase activity is necessary for the dynamic morphological extension of MG processes in the IPL. To this end, we first administered selective inhibitors of Mmp2 and Mmp9, ARP-100 and AG-L-66085, respectively, at 48 hpf, just prior to MG differentiation and the onset of their morphological elaboration. To assess the effect of removing Mmp2 or 9 activities from the onset of development on MG IPL process formation, we performed confocal imaging of MG membranes labelled by Tg(*tp1*:eGFP-CAAX) at 72 hpf (**Fig.3A**). We performed volumetric analysis of the processes of MG in the IPL and observed no differences between the treatments with either inhibitor compared to DMSO-treated controls (**Fig.3A-C**). To see whether blocking the activity of either enzyme later during MG process elaboration, following outgrowth initiation, we next administered the inhibitors from 72 hpf until 96 hpf (**Fig.3D**). Again, we observed no effect of Mmp2 or 9 inhibition on the overall complexity of MG processes (**Fig.3D-F**).

**Figure 3.**
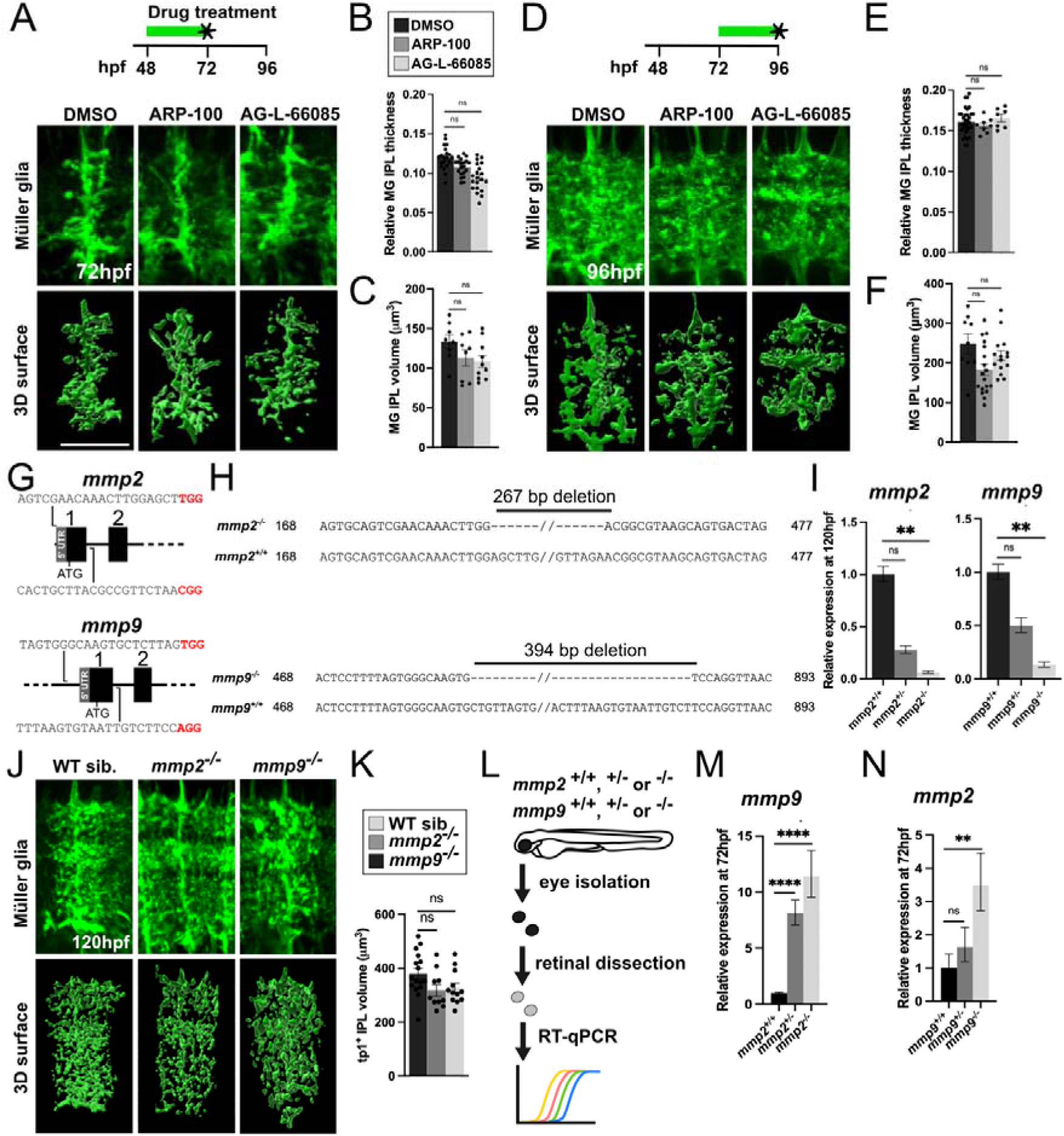
Inter-gelatinase redundancy and compensation mask phenotypes in loss of function *mmp2* and *mmp9* mutants. **(A)** Top: Pharmacological inhibition of Mmp2 (ARP-100) and Mmp9 (AG-L-66085) between 48 hpf and 72 hpf indicated by green line; time of imaging marked by a black star. Bottom: Confocal images of MG membrane protrusions in the IPL labelled by Tg(*tp1:eGFP-CAAX*) following treatment with inhibitors or DMSO at 72 hpf. Lower panels show the 3D surface reconstruction of MG processes. **(B)** Quantification of the thickness of MG IPL domain relative to retinal thickness. **(C)** Quantification of the MG IPL process volume following inhibitor treatment. **(D)** ARP-100 or AG-L-66085 treatment from 72 to 96 hpf; Confocal images of MG IPL processes at 96 hpf following inhibitor treatment**. (E)** Quantification of relative IPL thickness following Mmp2 or Mmp9 inhibition. (**F)** Quantification of MG process volume at 96 hpf following inhibitor treatment. **(G)** Exon 1 region targeted for CRISPR/Cas9 mutagenesis with two guide RNAs, generating an excision of the 5’UTR and exon 1 of the *mmp2* and *mmp9* genomic loci. **(H)** Alignment of sanger sequencing from homozygous *mmp2*^-/-^ and *mmp9^-/-^* mutants and WT DNA sequence, confirming a 270 and 394 bp-long deletion between the 2 gRNA binding sites for *mmp2* and *mmp9*, respectively. **(I)** Left: Relative expression of *mmp2* mRNA in *mmp2^-/-^, mmp2^+/-^* and WT siblings (*mmp2^+/+^)* whole larvae at 120 hpf; Right: Relative expression of *mmp9* mRNA in *mmp2^-/-^, mmp9^+/-^* and WT siblings (*mmp9^+/+^)* whole larvae at 120 hpf. **(J)** MG IPL processes labelled by Tg(tp1:eGFP-CAAX) in WT, *mmp2^-/-^*and *mmp9^-/-^* mutants at 120 hpf; bottom panels show 3D reconstruction of MG processes in the IPL. **(K)** Quantification of MG process volume in WT, *mmp2^-/-^* and *mmp9^-/-^* mutants at 120 hpf. **(L)** Summary of retinal isolation and RT-qPCR analysis. **(M-N)** Relative expression of *mmp9* in *mmp2* mutants and *mmp2* mRNA in *mmp9* mutant retinas compared to WT siblings and heterozygous mutants at 72 hpf. Scale bars, 10 µm.

To further investigate the consequences of Mmp2 or Mmp9 loss of function, we generated knockout mutants for both *mmp2* and *mmp9* using CRISPR-Cas9 mutagenesis. Both *mmp2* and *mmp9* genes contain only one paralogue in the zebrafish genome, and are located on chromosomes 7 and 8, respectively. We used two guide RNAs (gRNAs) to target exon 1 and the 5’ UTR or region upstream to excise the transcriptional start sites (TSSs) (**Fig.3G**). The excision of a 267 bp and 394 bp for *mmp2* and *mmp9*, respectively, were confirmed by PCR and subsequent Sanger sequencing and gel electrophoresis (**Fig.3G,H; Fig.S3A,B**). This was predicted to cause a frameshift and premature stop codon, resulting in null mutants. We compared two independently generated mutant alleles for each gene (**Fig.S4A,B**) and generated an additional mutation spanning exons 5 and 6 in *mmp2 (mmp2 ^e5/6^*^Δ*401*^) to produce a catalytically inactive mutant for comparison. There were no phenotypic differences observed between these mutants (**Fig.S4C**), and as such, we selected one TSS deletion allele for each gene for follow on analyses (*mmp2 ^e1^*^Δ*267*^ and *mmp2 ^e1^*^Δ*394*^). RT-qPCR validated a significantly reduced expression of the mRNA transcripts in heterozygous and homozygous mutants compared to WT sibling controls for both *mmp2* and *mmp9* mutants (**Fig.3I**). Additionally, western blot analysis was used to verify the loss of Mmp2 protein, as seen in the absence of bands corresponding to active and inactive Mmp2 (**Fig.S3C**). The Mmp9 antibodies tested did not result in appropriate bands in western blot, and hence we could not perform the same validation.

Consistent with the pharmacological inhibitors for Mmp2 and 9, our gross morphological analysis did not reveal any developmental defects in *mmp2^-/-^*and *mmp9^-/-^* mutant larvae compared to WT controls (**Fig.S3D**). The overall retinal organisation into laminae and cell densities between WT controls and mutants were also unchanged (**Fig.S3E-G**). Similarly, IF for common neuronal markers, HuC/D and PKCb which label ACs/RGCs, and BPCs, respectively, showed no discernible differences in neuronal morphology or numbers in homozygous mutants compared to WTs (**Fig.S3H-M**). IF for GS also revealed no effect on the overall MG morphology or numbers (**Fig.S3H,N**). Such lack of phenotypes was consistent across all mutant alleles generated (**Fig.S4**). We subsequently crossed the *mmp2* and *9* mutants with the MG membrane reporter Tg(*tp1:*eGFP-CAAX) fish to visualise MG processes in the IPL and performed imaging and volumetric analysis to compare their complexity in mutants and WT controls (**Fig.3J,K**). There was no difference in the volume of IPL processes between the genotypes, suggesting that MG processes develop normally in the absence of one gelatinase. To determine whether MG-neuron connections develop as normal in the *mmp2^-/-^* and *mmp9^-/-^* mutants, we next stained for the ribbon synapse marker Ribeye A and assessed the association between MG processes and Ribeye^+^ puncta (**Fig.S3O-R**). Both the number of Ribeye^+^ puncta and glia-synapse contacts were unchanged in mutants compared to WT controls (**Fig.S3Q,R**), further confirming that neuronal and glial development is unperturbed in the absence of Mmp2 or 9 alone.

Given the overlapping substrate specificities in the Mmp family, as well as potential genetic compensation mechanisms triggered by genetic knockouts in zebrafish, we asked whether the lack of phenotypes observed in our *mmp2 and 9* mutants could be due to transcriptional adaptation^53^. To this end, we performed RT-qPCR in retinas isolated from 72 hpf *mmp2* or *mmp9* mutants to assess the relative expression of the other gelatinase transcripts (**Fig.3L-N**). Interestingly, we found that *mmp2* expression increased 3.5-fold in *mmp9^-/-^* mutants compared to WT siblings, and in *mmp2^-/-^* mutants the relative expression of *mmp9* was approximately 12-fold higher than WTs (**Fig.3M,N**). Thus, functional redundancy between the gelatinases along with genetic compensation are likely to mask phenotypes in *mmp2^-/-^* and *mmp9^-/-^*mutants, akin to the lack of effect upon pharmacological inhibition of a single gelatinase (**Fig.3A-F**).

### Loss of both gelatinases results in reduced morphological complexity of MG processes and a disrupted ability to contact neuronal synapse

To overcome the genetic compensation in *mmp2^-/-^* and *mmp9^-/-^* mutants, we crossed these two mutants together to generate a pan-gelatinase (*mmp2^-/-^;mmp9^-/-^*) double mutant zebrafish. Gross morphological analysis at 120 hpf revealed a minor shortening of the anterior-posterior (A-P) larval body axis, as well as a reduction in the naso-temporal (N-T) length of the eye in the *mmp2^-/-^;mmp9^-/-^* mutants compared to WT controls (**Fig.4A-D**). These phenotypes were not apparent in the single mutants (**Fig.S3D-G**), pointing to a loss of the compensation observed in single gelatinase mutants. Given the decrease in mutant eye size, we subsequently examined global retinal organisation at 120 hpf and strikingly did not observe any gross morphological defects in retinal lamination in the *mmp2^-/-^;mmp9^-/-^* mutants, nor overall cell densities across the retinal layers, as reflected in their unchanged thicknesses (**Fig.S6A-C**). We next focused on the developing MG and assessed the number of GS^+^ MG cells at 120 hpf in the central retina, which was unchanged between *mmp2^-/-^;mmp9^-/-^*mutants and WTs (**Fig.S5D,E**). This indicates that gelatinases do not play a role in MG differentiation. Next, we assessed MG process morphology by crossing the double mutants with the MG membrane reporter, as before. Overall, MG processes in the IPL were not completely abolished in *mmp2^-/-^ ;mmp9^-/-^* mutants, however they did exhibit a reduced complexity compared to WTs (**Fig.4E**). The mean volume of MG processes in the IPL was decreased by 32% in the *mmp2^-/-^;mmp9^-/-^*mutants compared to WTs **(Fig.4E,F).** This indicates that gelatinase activity is required for MG process elaboration but not outgrowth initiation during MG morphogenesis and that the two gelatinases function redundantly in this context given the lack of phenotypes in the single mutants (**Fig.3;Fig.S4)**.

**Figure 4.**
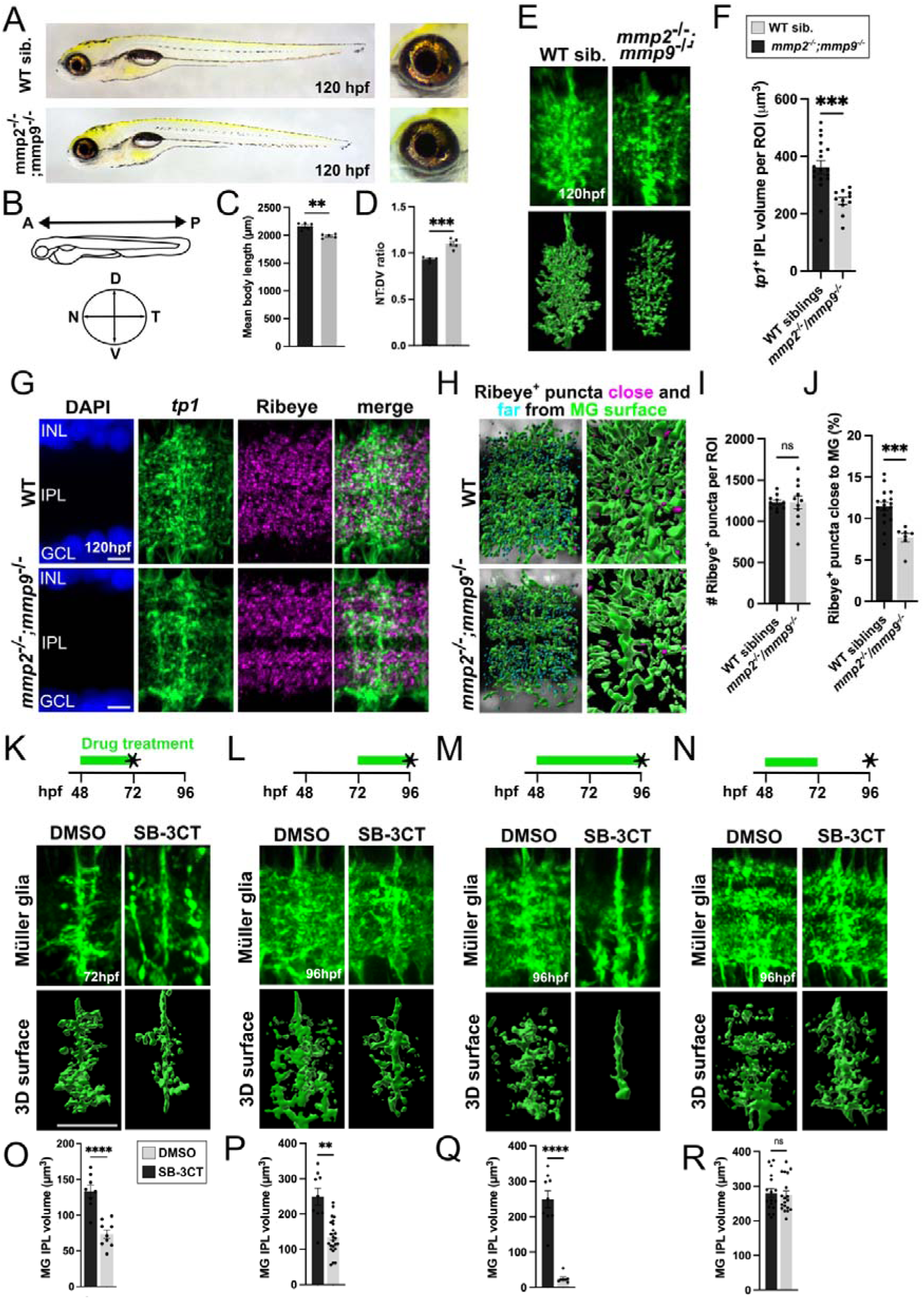
Mutants for *mmp2* and *mmp9* display healthy retinal development. **(A)** Whole mount images of WT and *mmp2*^-/-^/*mmp9*^-/-^ larvae at 120 hpf; right panels show close ups of the eyes. **(B)** Schematic of how body length and eye diameter measurements were performed; dorsal (D), ventral (V), nasal (N) and temporal (T). **(C)** Quantification of body axis length in WT and *mmp2*^-/-^;*mmp9*^-/-^ mutants at 120 hpf. **(D)** Quantification of the ratio of the D-V and N-T diameter measurements in WT and *mmp2*^-/-^;*mmp9*^-/-^ mutants. **(E)** MG IPL processes at 120 hpf labelled by Tg(*tp1*:EGFP-CAAX); Lower panels show a 3D surface reconstruction of membrane-GFP tagged MG cell processes. **(F)** Quantification of MG process volume in the IPL of WT and *mmp2*^-/-^;*mmp9*^-/-^ mutants at 120 hpf. **(G)** IF for RibeyeA to label ribbon synapses in the IPL of WT and *mmp2^-/-^*;*mmp9^-/-^*retinas at 120 hpf. MG membranes labelled in green and nuclei in blue (DAPI). **(H)** Reconstruction of RibeyeA synaptic puncta far (cyan) and close (magenta) to MG process surfaces (green) in double mutants compared to WTs. **(I)** Quantification of the number of Ribeye+ spots in in the IPL per ROI between genotypes. **(J)** Quantification of the number of Ribeye+ spots close to MG surfaces in WT and *mmp2*^-/-^;*mmp9*^-/-^ mutants at 120 hpf. **(K)** Retinas of Tg(tp1:eGFP-CAAX) larvae treated with DMSO or 10µM SB-3CT from 48-72 hpf (green bar). Black asterisk denotes the time of retinal imaging. **(L)** Inhibitor treatment from 72-96 hpf. **(M)** Inhibitor treatment from 48-96 hpf. **(N)** Inhibitor treatment from 48-72 hpf, followed by removal of inhibitors and imaging at 96 hpf (black asterisk). **(O-R)** Quantification of the volume of MG processes in the IPL following the SB-3CT inhibitor treatments in K-N. Scale bars, 10µm.

Having observed that MG processes in the IPL do not elaborate to the same degree in *mmp2^-/-^;mmp9^-/-^*mutants, we next sought to explore how this impacts their ability to contact neuronal synapses at 120 hpf, a point at which retinal circuitry is largely established. As before, we performed antibody staining for Ribeye A and assessed the proportion of synaptic puncta in close proximity to MG processes **(Fig.4G-J)**. Firstly, the mean number of Ribeye^+^ synaptic puncta was unchanged in the pan-gelatinase mutants compared to WTs **(Fig. 4I)**, indicating that synaptogenesis itself was unaffected in the *mmp2^-/-^;mmp9^-/-^*mutants. Since the volume of MG processes was reduced, they occupied less space in the IPL and associated with Ribeye puncta to a significantly reduced degree in comparison to WT retinas **(Fig.4H,J)**. We observed a similar decrease in the percentage of Ribeye puncta close to MG processes in *mmp2^-/-^;mmp9^-/-^*mutants compared to WTs **(Fig.4J)**. Furthermore, we also used a known pan-gelatinase inhibitor, SB-3CT, to simultaneously block Mmp2 and Mmp9 activities with precise temporal control to validate the effect of gelatinase inhibition on MG development. We first applied the inhibitor at 48 hpf to eliminate gelatinase activity from the onset of MG development (**Fig.4K**) and assessed MG process volume in the IPL 24 hours after treatment. Similarly to the *mmp2^-/-^;mmp9^-/-^*mutants, we observed a reduced complexity of MG processes in the IPL following treatment with SB-3CT, as revealed in the 50% reduction in their volume compared to DMSO-treated controls (**Fig.4K,L**). Since neuronal development is also still taking place at 48 hpf when the inhibitor was administered^62^, we could not exclude that gelatinase inhibition may influence neurons and therefore elicit secondary effects on MG morphology. To circumvent any possible secondary effects upon MG morphogenesis via neuronal disruption, we also performed the same experiment at 72 hpf, once neuronal differentiation is largely complete and most of the neuronal connections have been established^63^ (**Fig.4M**). SB-3CT treatment from 72 to 96 hpf resulted in a similar 46% reduction in MG process volume compared to DMSO controls (**Fig.4N**). Further, we found that a prolonged exposure to SB-3CT from 48 to 96 hpf greatly exacerbated the reduction in MG morphological complexity, as reflected in a 90% reduction in MG IPL volume compared to DMSO-controls (**Fig.4O,P**). On the other hand, removal of the inhibitor after an initial exposure from 48 to 72 hpf led to a restoration of MG IPL volume to the same levels as DMSO controls by 96 hpf (**Fig.4Q,R**). Thus, simultaneous removal of Mmp2 and Mmp9 activity impairs the outgrowth and elaboration of MG processes, and can be reversed by alleviating gelatinase inhibition, confirming that gelatinase activity is required throughout MG morphogenesis. Moreover, SB-3CT treatment was found to reduce contacts between Ribeye^+^ synaptic puncta and MG processes by 50% without impairing synaptic numbers (**Fig.S5**), akin to our observations in the *mmp2^-/-^;mmp9^-/-^*mutants, while single-gelatinase inhibition had no effects.

Together, these findings show that gelatinase activity is important for the morphological elaboration of MG processes and their integration into neuronal circuits, and that Mmp2 and 9 function redundantly in this context alongside genetically compensating for each other’s absence.

### Gelatinase-deficient Müller glia display signs of gliosis in early adulthood

As MG are the principal support cell in the retina, express the ECM modifying gelatinases, and show defects in the formation of glia-synapse contacts in their absence during development, we hypothesised that there would be premature signs of degeneration in the retina of gelatinase mutants due to impaired glial support. First, as we showed that MG morphology is not established to the same degree of complexity in the *mmp2^-/-^;mmp9^-/-^* mutants compared to WTs, we assessed whether this persists in the mature retina post-larval development by performing IF for GS (**Fig.5A**). We first assessed MG numbers at 3- and 6-months post fertilisation (mpf) and found that the number of MG cells was unaffected at 3 mpf, while at 6 mpf, we saw a subtle but significant reduction in MG cell numbers in the *mmp2^-/-^;mmp9^-/-^*mutants **(Fig.5B,C)**. We also confirmed that the volume of GS^+^ MG IPL processes was still significantly lower in *mmp2^-/-^;mmp9^-/-^* mutants compared to WTs at 6 mpf (**Fig.5D-E**), indicating a sustained reduction in MG morphological complexity in the absence of gelatinases throughout life. This reduced IPL process complexity contrasted with the rest of the cell, which showed signs of MG hypertrophy at 6 mpf, as reflected in the visible expansion and morphological disorganisation of the GS^+^ cells in the other morphological compartments, such as the cell bodies (**Fig.5A,A’**). Since glial hypertrophy is a hallmark of gliosis, we next investigated the expression of the glia-specific intermediate filament protein Glial fibrillary acidic protein (Gfap), which is not only a structural marker of MG but also a proxy for stress and gliotic activation (**Fig.5G**) ^64–66^. At 3 mpf, Gfap immunoreactivity showed the characteristic localisation in the basal MG end feet in both WTs and mutants. At 6 mpf however, we observed Gfap staining in more apical regions of the cells, again indicative of gliosis (**Fig.5G**). 3D segmentation and quantification of the volume of Gfap staining in apical vs basal regions confirmed these observations and showed a significant increase in apical: basal Gfap distribution at 6 mpf, but not 3 mpf (**Fig.5H-J**).

**Figure 5.**
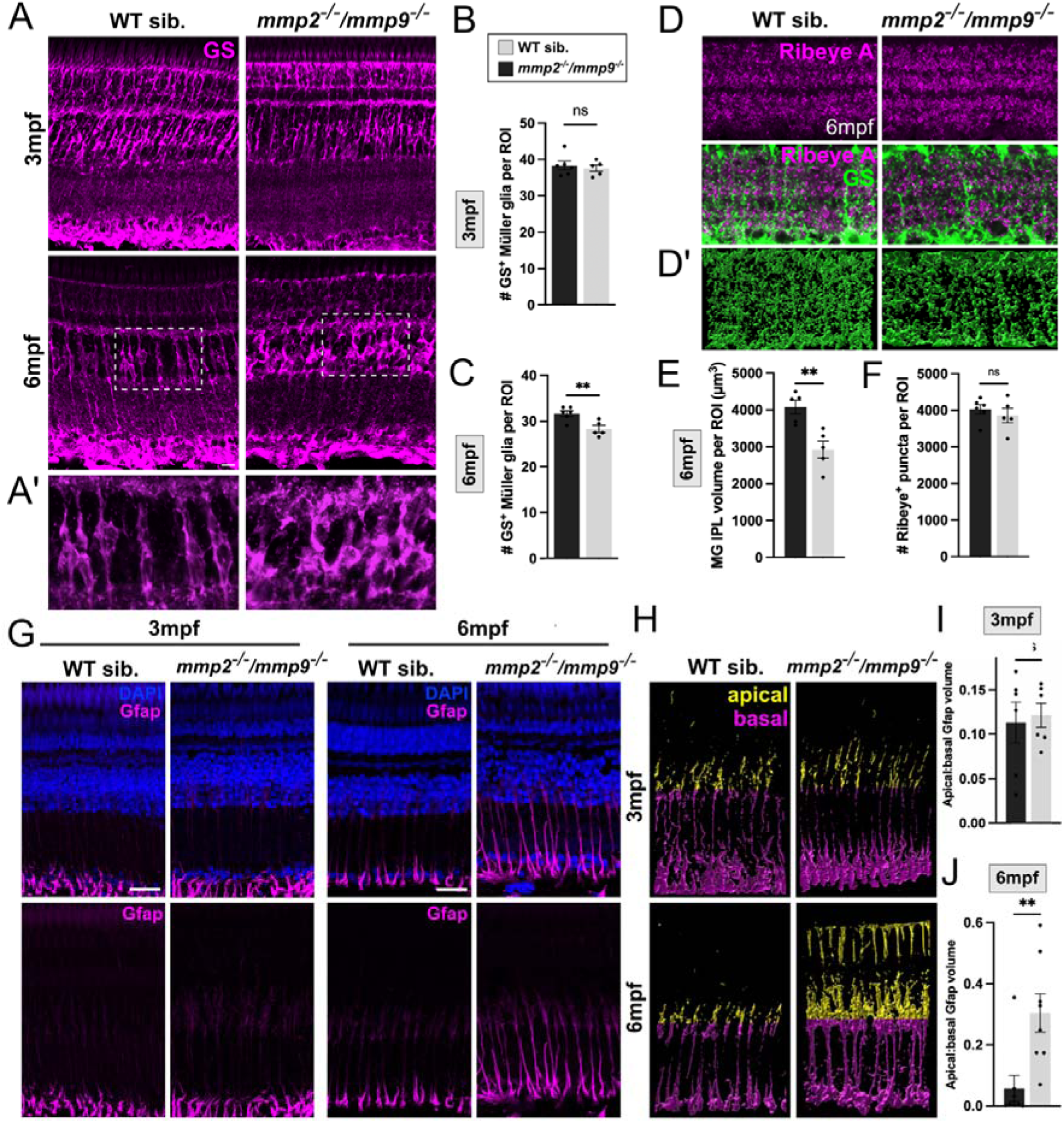
Müller glia display morphological signs of gliosis in the absence of the gelatinases in the adult retina. **(A)** IF for glutamine synthetase (GS) in retinal cryosections at 3- and 6-months post fertilisation (mpf). Scale bars, 10 µm. **(A’)** Zoom box showing MG hypertrophy in the inner nuclear layer. **(B)** Quantification of the number of GS-positive MG per region of interest (ROI) at 3 mpf in wild type (WT) siblings and *mmp2^-/-^;mmp9^-/-^* mutants. **(C)** Quantification of the number of GS-positive Müller glia per ROI at 6 mpf in wild type (WT) siblings and *mmp2^-/-^;mmp9^-/-^*mutants. **(D)** Immunostaining for Ribeye A (magenta) and GS (green) in retinal cryosections from WTs and *mmp2^-/-^;mmp9^-/-^* mutants. **(D’)** 3D surface reconstruction of Ribeye and GS immunostaining **(E)** Quantification of the MG volume of IPL processes per ROI at 6 mpf in WT siblings and *mmp2^-/-^;mmp9^-/-^*mutants. (F) Quantification of the number of MG volume of IPL processes per ROI at 6 mpf in WT siblings and *mmp2^-/-^;mmp9^-/-^*mutants. (G) Antibody staining for Gfap (magenta), nuclei labelled with DAPI (blue) in WT siblings and *mmp2^-/-^;mmp9^-/-^* mutants at 3 and 6 mpf. **(H)** 3D surface reconstruction of Gfap antibody staining in G, divided into apical and basal halves for quantification; yellow=apical, magenta=basal. **(I)** Comparison of the ratio of apical:basal Gfap volume in 3D surface reconstructions from H at 3 mpf. **(J)** Comparison of the ratio of apical:basal Gfap volume in 3D surface reconstructions from H at 6 mpf.

These findings reveal that the reduced MG IPL process complexity in *mmp2^-/-^;mmp9^- /-^* mutants persists into adulthood, and by 6 mpf is coupled with morphological signs of gliotic activation, pointing to potential neurodegeneration in the *mmp2^-/-^;mmp9^-/-^*mutants.

### *mmp2^-/-^;mmp9^-/-^* mutants exhibit neuronal degeneration and declining visual activity with age

In addition to a sustained reduction in MG process complexity and gliotic activation, we also assessed the retinal immune cells, the microglia, by IF at 6 mpf (**Fig.6A**). The overall numbers of microglia were unchanged between the WTs and *mmp2^-/-^ ;mmp9^-/-^* mutants, as assessed with markers Lcp1^+^ and 4c4^+^ (**Fig.6B,C**). However, when assessing their localisation across the retinal layers, we noticed a tendency to occupy more apical layers, such as the ONL and photoreceptor layer (PRL), when assessed by Lcp1 staining (**Fig.6D**), and a similar, but statistically significant shift of 4c4^+^ cells towards the outer retina (**Fig.6E**). Finally, we assessed the morphology of the microglial cells and observed an increased incidence of activated microglia characterised by their amoeboid-like morphology, compared to the WTs, which had mostly ramified microglia (**Fig.6A’**). Quantification of the number of ramified vs amoeboid microglia revealed a significant increase in amoeboid cells and a reciprocal decrease in those with a ramified morphology in *mmp2^-/-^;mmp9^-/-^* mutants compared with WT siblings.

**Figure 6.**
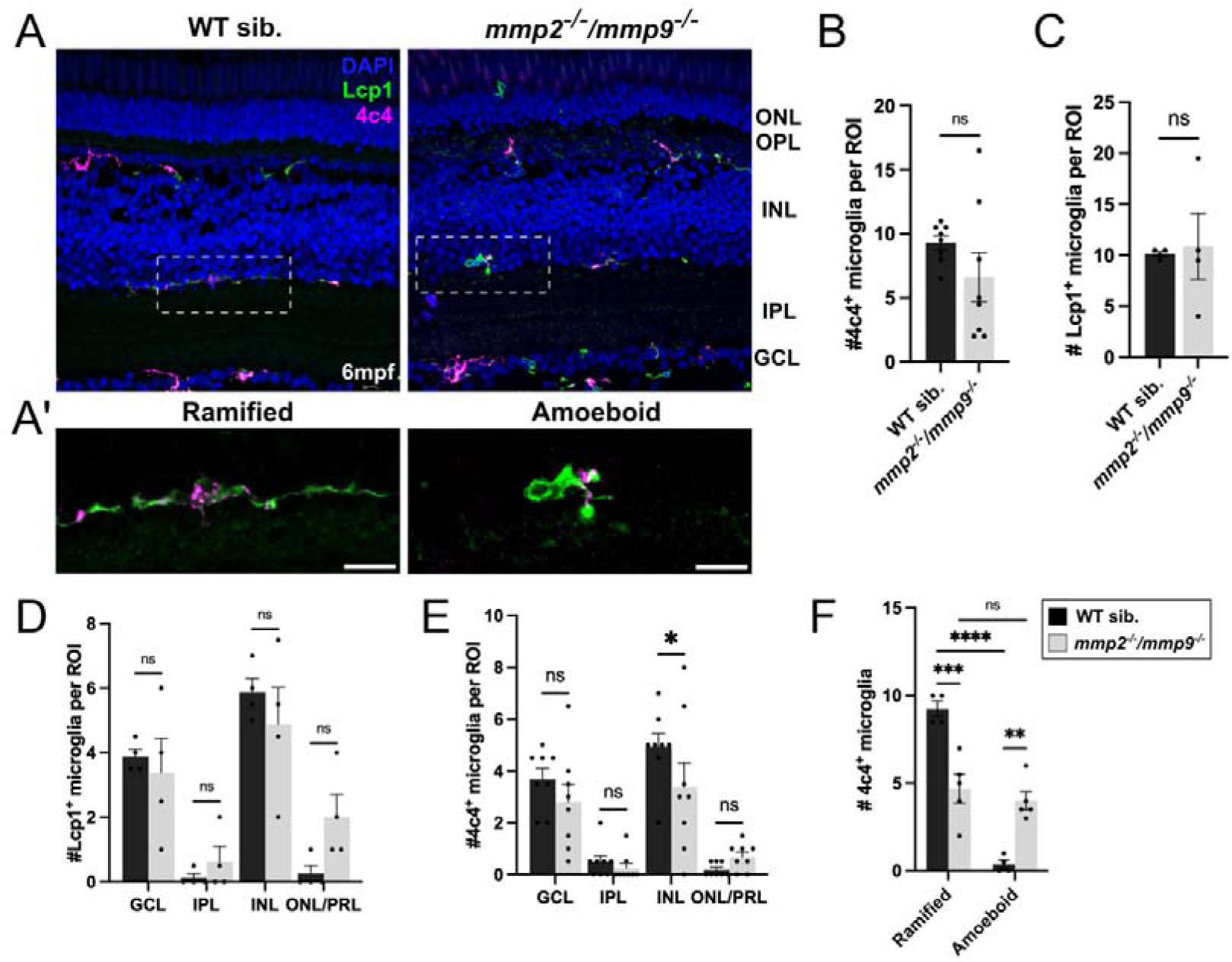
Microglial activation in the retina of pan-gelatinase mutants. **(A)** Immunostaining for microglial markers L-plastin (Lcp1; green) and 4c4 (magenta) in retinal cryosections at 6-months post fertilisation (mpf). Nuclei are labelled with DAPI (blue). **(A’)** Zoom boxes showing ramified microglia in wild-type (WT) siblings and amoeboid (activated) microglia in *mmp2^-/-^/mmp9^-/-^*mutants. **(B)** Quantification of the number of 4c4-positive microglia per 100µm x 100µm x 10µm region of interest (ROI) at 6 mpf in WT siblings and *mmp2^-/-^/mmp9^-/-^* mutants. **(C)** Quantification of the number of Lcp1-positive microglia per 100µm x 100µm x 10µm region of interest (ROI) at 6 mpf in WT siblings and *mmp2^-/-^/mmp9^-/-^* mutants. **(D)** Quantification of the number of Lcp1-positive microglia per 100µm x 100µm x 10µm ROI at 6 mpf in WT siblings and *mmp2^-/-^/mmp9^-/-^* mutants. **(E)** Quantification of the number of 4c4-positive microglia per 100µm x 100µm x 10µm ROI at 6 mpf in WT siblings and *mmp2^-/-^/mmp9^-/-^*mutants; GCL. **(F)** Quantification of the number of ramified and amoeboid microglia labelled with 4c4 antibody per 100µm x 100µm x 10µm ROI at 6 mpf in WT siblings and *mmp2^-/-^/mmp9^-/-^* mutants. Scale bars, 10µm.

Given the gliosis-like phenotypes observed in adult MG in *mmp2^-/-^;mmp9^-/-^* mutants, we next assessed whether neuronal cells were also affected by the loss of both gelatinases. To this end, we compared the gross morphology and numbers of different neuronal types by IF from development to adulthood. We observed no differences in the number and overall appearance of the major neuronal classes in the retina ACs, RGCs, PBCs or PRs in the *mmp2^-/-^;mmp9^-/-^*mutants compared to WTs at 120 hpf or 3 mpf (**Fig.7A-O**), labelled by HuC/D, PKCb and Gnat2, respectively. At 6 mpf however, there was a significant reduction in the numbers of all cell types, pointing to a global neurodegeneration in *mmp2^-/-^;mmp9^-/-^* mutants **(Fig.7E, H,K,O),** consistent with the gliosis-like response observed among the glial cells. Given the reduced numbers of neurons observed in *mmp2^-/-^;mmp9^-/-^* mutants, we next sought to assess how this may impact visual function of the pan-gelatinase mutants and WTs. We performed optokinetic response (OKR) analysis to assess the visual responses of larvae at 120 hpf and adult fish at both 3 and 6 mpf **(Fig.7P-Q)**. Interestingly, at 120 hpf when robust vision commences to support the emergence of prey capture behaviour^63^, there was no difference in the number of eye saccades per minute in *mmp2^-/-^;mmp9^-/-^* mutants compared to WT siblings, under varying spatial frequency, grating velocity or contrast of the visual stimulus **(Fig.7P,Q)**. The number of eye saccades per minute was also unaffected by the absence of gelatinases at 3 mpf, suggesting that visual function was unaffected in the mutants during early adulthood, when the fish reach sexual maturity **(Fig.7Q)**. However, by 6 mpf, we observed a significant decline in eye saccades in response to the visual stimulus in the pan-gelatinase mutants compared to WTs (**Fig.7Q**), coinciding with the apparent retinal degeneration occurring at this time.

**Figure 7.**
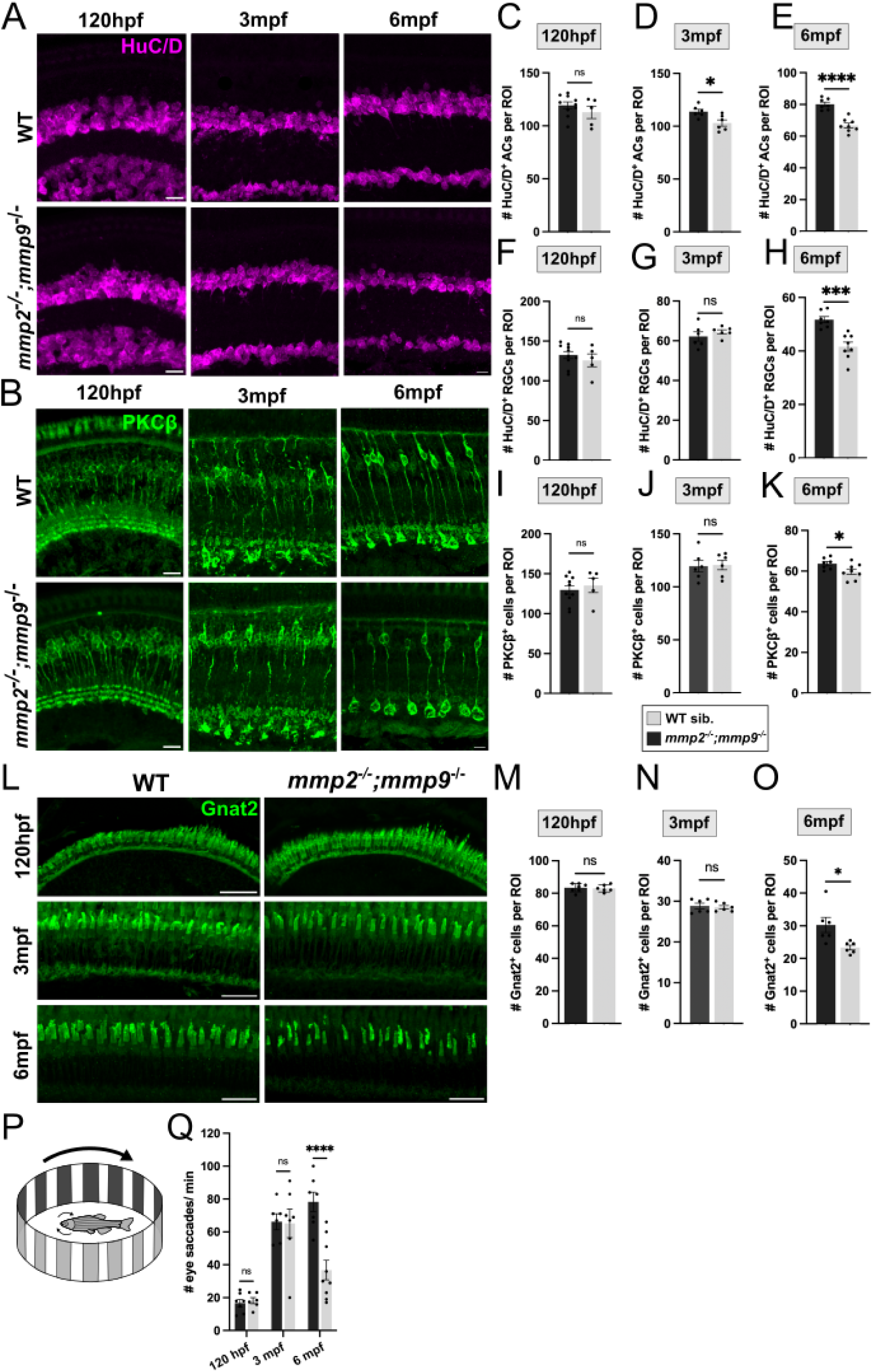
Neurodegeneration and visual function defects in adult *mmp2^-/-^ ;mmp9^-/-^* mutants. **(A)** Immunostaining for neuronal marker HuC/D to label ACs and RGCs in retinal cryosections at 120 hpf, 3 mpf and 6 mpf, in WT and *mmp2^-/-^;mmp9^- /-^* mutants. **(B)** Immunostaining for BPC marker PKCβ in WT and *mmp2^-/-^;mmp9^-/-^*mutants at 120 hpf, 3 mpf and 6 mpf. **(C-E)** Quantification of the number of HuC/D-positive ACs in a 100x100x10 µm ROI at 120 hpf, 3 mpf and 6 mpf in WT and pan-gelatinase mutants. **(F-H)** Quantification of the number of HuC/D-positive RGCs in a 100x100x10µm ROI at 120 hpf, 3 mpf and 6 mpf in WT and *mmp2^-/-^;mmp9^-/-^*mutants. **(I-K)** Quantification of the number of PKCβ-positive BPCs in a 100x100x10µm ROI at 120 hpf, 3 mpf and 6 mpf in WT and *mmp2^-/-^;mmp9^-/-^*mutants. **(L)** Immunostaining for cone photoreceptor marker Gnat2 in WT and *mmp2^- /-^;mmp9^-/-^* mutants at 120 hpf, 3 mpf and 6 mpf. **(M-O)** Quantification of the number of Gnat2-positive photoreceptor cells in a 100x100x10µm ROI at 120 hpf, 3 mpf and 6 mpf in WT and *mmp2^-/-^;mmp9^-/-^*mutants. **(P)** Schematic of adult OKR experimental set up. **(Q)** Comparison of mean number of eye saccades per minute (average of both eyes) at 120 hpf, 3 mpf and 6 mpf. Scale bars, 10µm.

Taken together, these findings indicate that while initially neuronal development and visual function are unperturbed in the absence of gelatinases, the improper establishment of glial support in the synaptic layer results in the eventual neurodegeneration, inflammation and visual dysfunction in the adult retina.

## Discussion

Our findings identify a central role for the gelatinases Mmp2 and Mmp9 in shaping Müller glial (MG) morphology within the zebrafish retina. Combined loss of both enzymes, achieved through pharmacological inhibition or genetic ablation, caused a substantial reduction in the volume and complexity of MG processes in the IPL. This was found to impair their ability to associate with neuronal synapses, a key interaction that supports effective neurotransmission ^67,68^. These defects persisted into adulthood, demonstrating that gelatinase activity is necessary not only for the initial elaboration of MG arbours during development but also for maintaining their structure and synaptic associations throughout life.

Early in development, the effects of gelatinase loss were confined to MG, with no measurable impact on neuronal number or function at 120 hpf. By 6 mpf, however, we observed reduced visual performance, progressive neuronal loss, and widespread gliotic activation. The appearance of MG morphological defects prior to neuronal decline suggests that impaired glial support contributes indirectly to the neurodegeneration observed later in life. The loss of fine MG protrusions in the IPL, critical for forming tripartite synapses and maintaining neurotransmitter homeostasis, likely compromises synaptic physiology within this layer. Since MG processes were not completely abolished, partial preservation of glial-neuronal contacts likely delayed the onset of neuronal dysfunction. Interestingly, AC numbers began to decline from 3 mpf, much earlier than other cell types, suggesting either a heightened vulnerability to the effects of gelatinase loss or a particularly prominent reliance on MG support. Although our analysis focused on the IPL, MG span all retinal layers and associate with multiple ECM-rich domains, including the OPL, interphotoreceptor matrix ^69^, and limiting membranes. Disruption of Mmp2/9 function may therefore perturb additional MG compartments, and thereby contribute to the global neuronal degeneration observed, including in PRs that do not directly synapse within the IPL. It is likely that MG process outgrowth is regulated by parallel mechanisms and gelatinases play an analogous role in the OPL, and perhaps even other CNS neuropils. Likewise, reduced interactions between MG end feet and the vasculature may further disturb the neuro-glio-vascular unit^70^, providing additional routes by which gelatinase loss could compromise retinal homeostasis.

While we know that gelatinases are important for proper outgrowth and elaboration of MG processes, it remains unclear which specific Mmp2/9 substrates are important in this context. Since MMP targets are widespread and extend beyond the ECM^32,33,71–73^, it will be important to identify those specifically required for MG process elaboration. While ECM components remain the obvious candidates, MMPs have also been shown to degrade non-ECM proteins localised on the cell membrane, such as cadherins or integrins, or those secreted or bound to the ECM, like the growth Tgf-β which is known to bind Decorin, an MMP substrate^74^. In addition to their canonical extracellular functions, MMPs have also been described to act intracellularly, and their expression has been detected in the cytosol and within organelles like the nucleus and mitochondria in a range of cell types, including MG^75–81^. MMP2 has been shown to be inefficiently secreted out of cells, with approximately half of the nascent protein retained inside cells, which is thought to be an evolutionarily conserved effort to ensure their intracellular functions^77,81^. Among intracellular targets of MMPs are proteins relating to the cytoskeleton, such as actin, ARP2/3, intermediate filament proteins Vimentin, Desmin and Stathmin^82^, all of which could alter the outgrowth of protrusions and are hence possible candidate substrates in MG development. Future proteomic and degradomic approaches will be essential for identifying the key gelatinase substrates that drive MG process elaboration in the retina.

Moreover, despite their reported ubiquitous expression and diverse roles across virtually all organs and tissues, we observed a lack of phenotypes following the loss of a single gelatinase either by pharmacological inhibition or its genetic deletion. Only the simultaneous deletion of both gelatinases led to more noticeable phenotypes in the retina and systemically in the rest of the body. This raises an important consideration in MMP research, which is the redundancy across this enzyme family alongside possible inter-MMP genetic compensation and transcriptional regulation that may occur. This was not surprising given the subtlety of phenotypes in the MMP2 knockout mice ^83,84^, which was also attributed to the upregulation of MMP9^85^. While the loss of a single MMP may have minimal effects on physiological processes, this may differ in the context of disease or stress on a particular system, which has been documented across different disease and injury models in MMP-deficient backgrounds. Any characterisation of MMP9 null mutants is scarce, likely due to a lack of obvious phenotypes, and indeed most studies have instead explored the consequences of its loss in models of neurodegeneration or ischaemia ^86–89^. In the retina, the loss of Mmp9 was also only explored in the adult retina following damage, where the absence of the gelatinase led to an increased inflammatory response, proliferation and PR regeneration^41,90^.

Finally, MMP2/9 expression has been described in most types of glia, including astrocytes, oligodendrocytes and microglia^91–93^, pointing to a potentially conserved requirement for these enzymes across all glia. Astrocytes, the brain analogue of the MG, have been shown to express MMP2 and 9 specifically at their fine filopodial processes^94,95^, and exhibit morphological defects in their absence, further suggesting that the novel role of gelatinases in MG process outgrowth identified in the current study may be relevant beyond the zebrafish retina. Additionally, ECM remodelling and MMP dysregulation are known hallmarks of CNS pathologies^72^ and coincide with gliosis-associated morphological changes^96–103^. Thus, understanding how glia may utilise enzymes such as gelatinases to help establish their precise positions and elaborate morphologies in busy extracellular environments and maintain them throughout life, will be essential to unravelling the mechanisms of glial dysregulation in disease, which in turn impact neuronal health. Preventing these pathological changes may pose novel considerations for preventing or potentially halting retinal degeneration via the support cells, particularly in light of our findings that glial cell dysfunction can precede neurodegeneration. This is especially important considering that gliosis occurs in virtually all neurodegenerative diseases, injury models and during aging^5,32,65,72,104–106^, and thus represents an appealing universal therapeutic target for intervention.

## Conclusions

This work establishes a critical role for Mmp2/9 in regulating MG morphology, with implications for retinal development, maintenance, and degeneration. We identify a novel function for gelatinase-mediated ECM remodelling in MG process elaboration, linking glial morphology to neuronal health. More broadly, these findings highlight conserved mechanisms by which glia shape and sustain neural circuits, offering potential avenues for therapeutic intervention in neurodegenerative diseases. Promoting or preserving glial support may represent an effective strategy for mitigating neuronal degeneration across the CNS.

## Acknowledgments

We thank Bryan Crawford for generous sharing of embryos from the Tg(*hsp70:*EMMA-Mmp2) line and assistance with analysis. We also thank Mattias Loidolt and Isaac Bianco for assistance with OKR measurements. We thank Nicole Noel and Yi Jiang for critical reading of the manuscript. This work was supported by Moorfields Eye Charity for funding to develop the Institute of Ophthalmology zebrafish system (GR001114) and a PhD studentship to RBM (GR001148). This work was supported by a BBSRC David Phillips Fellowship (BB/S010386/1) and BBSRC grant (UKRI705) to RBM.

