## Supplemental Figures for "Gelatinase activity is required for Müller glia morphogenesis and neuronal maintenance in the retina"

**SUPPLEMENTARY FIGURES AND TABLES**

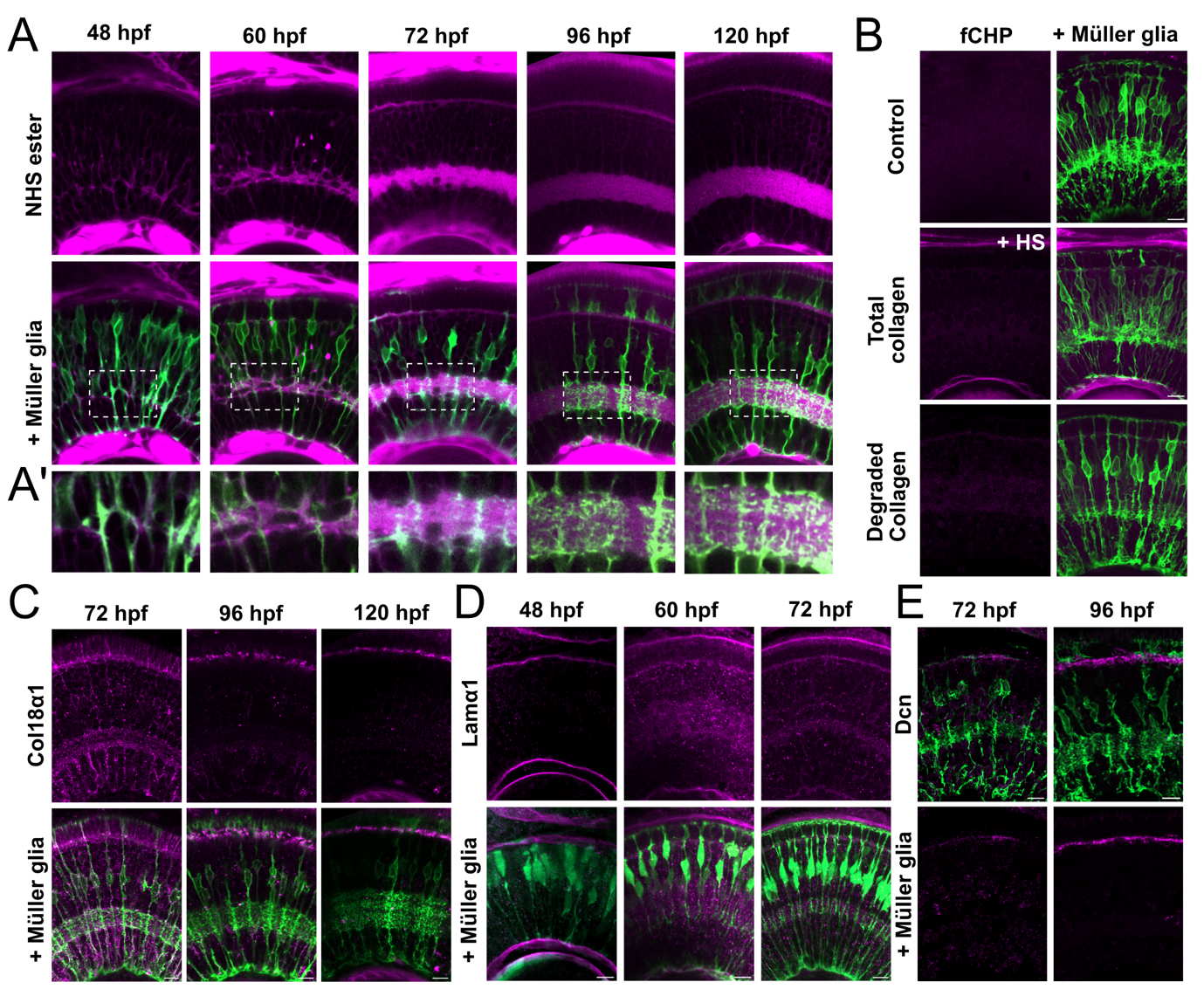

**Figure S1. ECM landscape during retinal development. (A)** NHS ester-647 (magenta) injected retinas throughout MG morphogenesis (48-120 hpf); Müller glia (green) labelled by Tg(tp1:eGFP-CAAX).**(A’)** Close up of NHS ester staining in the IPL. **(B)** fluorescent collagen hybridising peptide (fCHP) staining in the retina at 72 hpf; Controls incubated with inactive fCHP, total collagen labelled by heat shock of the larvae to induce collagen degradation, degraded collagen labelled in non-heat shocked larvae. **(C)** Col18a1 (magenta) antibody staining in the retina at 72, 96 and 120 hpf. Müller glia (green) labelled by Tg(tp1:eGFP-CAAX); Top panels show single z-planes, bottom panels show maximum intensity z-projections. **(D)** Lama1 (magenta) antibody staining in the retina at 48, 60 and 72 hpf. Müller glia (green) labelled by Tg(tp1:eGFP-CAAX); Top panels show single z-planes, bottom panels show maximum intensity z-projections. **(C)** Col18a1 (magenta) antibody staining in the retina at 72 and 96 hpf. Müller glia (green) labelled by Tg(tp1:eGFP-CAAX); Top panels show single z-planes, bottom panels show maximum intensity z projections. Scale bars, 10µm.

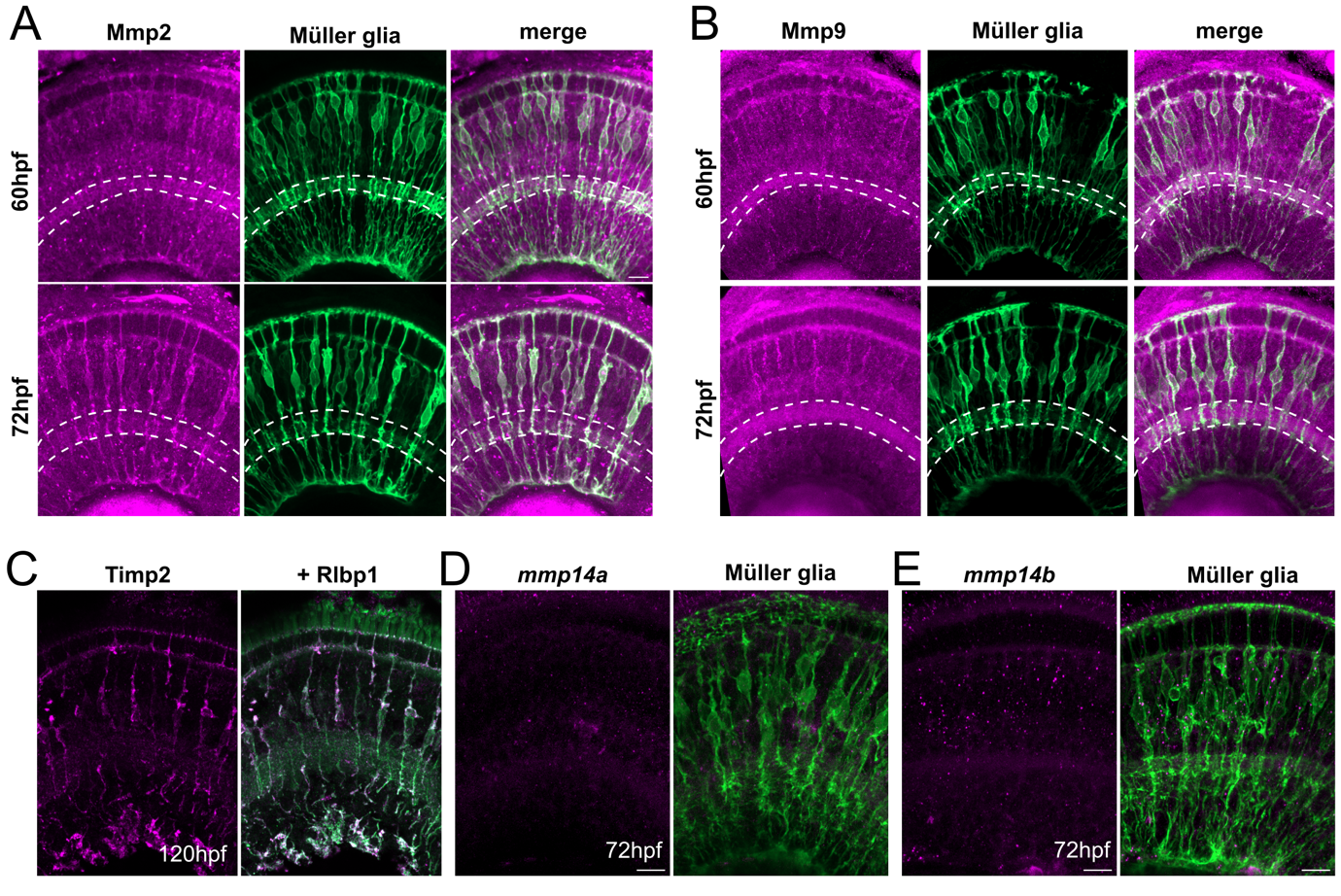

**Figure S2. Mmp and Timp expression patterns during retinal development. (A)** IF for Mmp2 (magenta) in Tg(tp1:eGFP-CAAX) larvae labelling Müller glia cells in the zebrafish retina. Dotted lines denote the synaptic inner plexiform layer (IPL) at 60 and 72 hpf. **(B)** IF staining for Mmp9 at 60 and 72 hpf. **(C)** Timp2 (magenta) antibody staining on 120 hpf zebrafish retinal cryosections co stained with anti-Rlbp1 (green), labelling Müller glia. **(D)** HCR-FISH detection of *mmp14a* mRNA in Tg(tp1:eGFP-CAAX) retinas at 72 hpf. **(E)** HCR-FISH detection of *mmp14b* mRNA in Tg(tp1:eGFP-CAAX) retinas at 72 hpf. Scale bars, 10 µm.

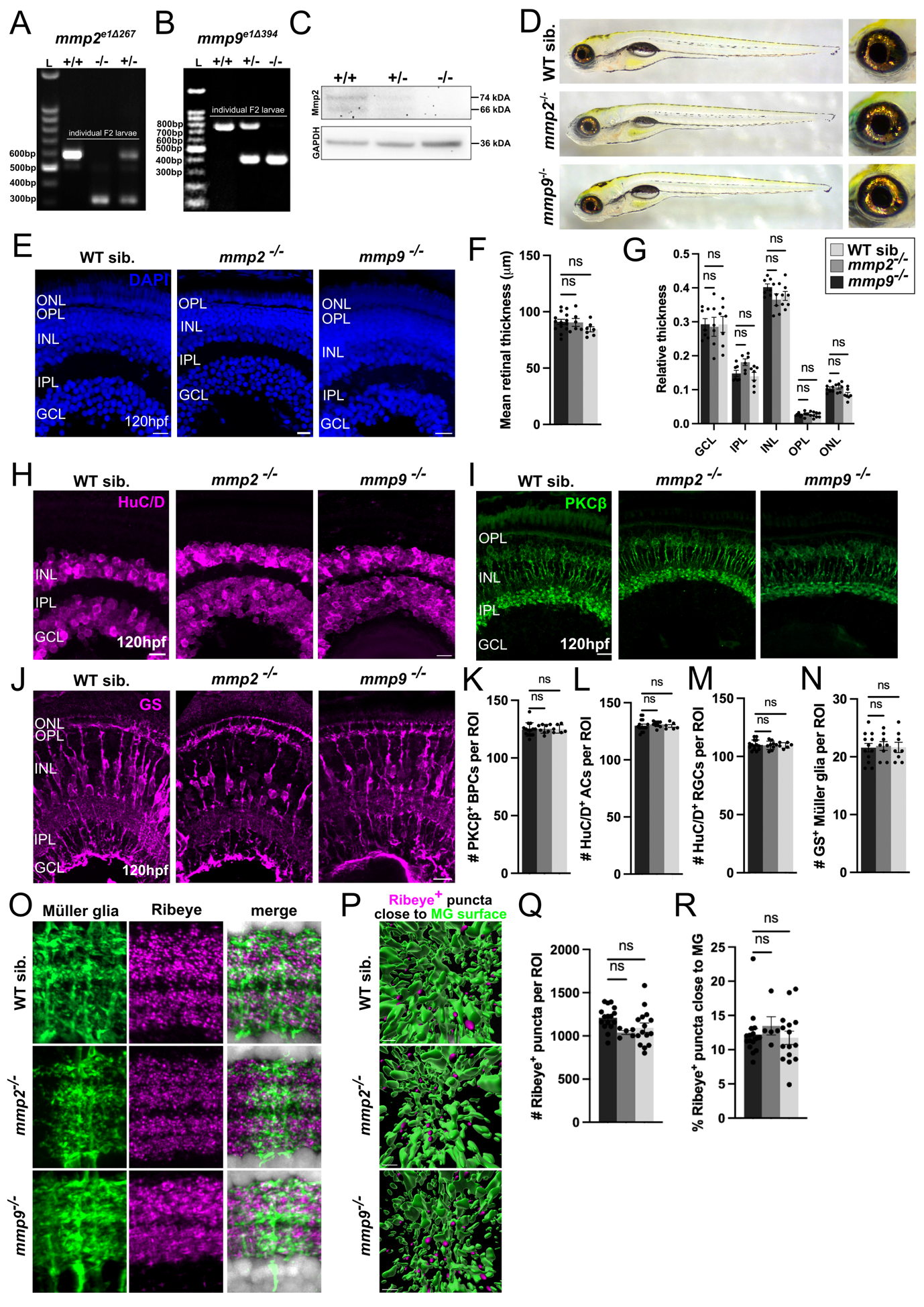
**Figure S3. Loss of function of Mmp2 or Mmp9 alone has no effect on retinal development. (A)** Gel electrophoresis showing region flanking the CRISPR/Cas9 deletion amplified from genomic DNA of *mmp2* homozygotes (+/+), heterozygotes (+/-) and WT siblings (+/+) from the F2 generation. **(B)** Gel electrophoresis showing the genomic deletion of *mmp9* fragment in F2 generation- larvae*.* **(C)** Western blot detection of zebrafish Mmp2 in pooled whole-larval lysates at 120 hpf. **(D)** Whole mount images of *mmp2^-/-^* or *mmp9*^-/-^ mutant and WT larvae at 120 hpf. Right panels show a zoom of the eyes. **(E)** Overall retinal lamination visualised by nuclear DAPI staining in 120 hpf-old *mmp2^-/-^* , *mmp9*^-/-^ mutant or WT larvae. **(F)** Quantification of apicobasal retinal thickness from images in E. and **(G)** the relative thicknesses of each retinal layer compared to overall retinal thickness. **(H)** Amacrine and retinal ganglion cell labelled by HuC/D in 120 hpf-old *mmp2^-/-^* , *mmp9*^-/-^ mutant or WT retinas at 120 hpf. **(I)** Anti-PKCβ staining labelling bipolar cells across the genotypes. **(J)** MG labelled by anti-GS antibody at 120 hpf. **(K-N)** Quantification of cell numbers for each retinal cell type mentioned in H-J. **(O)** IF for RibeyeA (magenta) in mutant and WT retinas at 120 hpf; Nuclei labelled with DAPI (grey), MG membranes labelled by Tg(*tp1*:eGFP-CAAX) (green). **(P)** 3D reconstruction of MG processes (green) and Ribeye+ puncta (magenta spots) which were defined as close to MG surfaces. **(Q)** Quantification of the mean number of Ribeye^+^ puncta for ROI, as shown in O. **(R)** Quantification of percentage of Ribeye^+^ spots close to MG processes in the IPL.

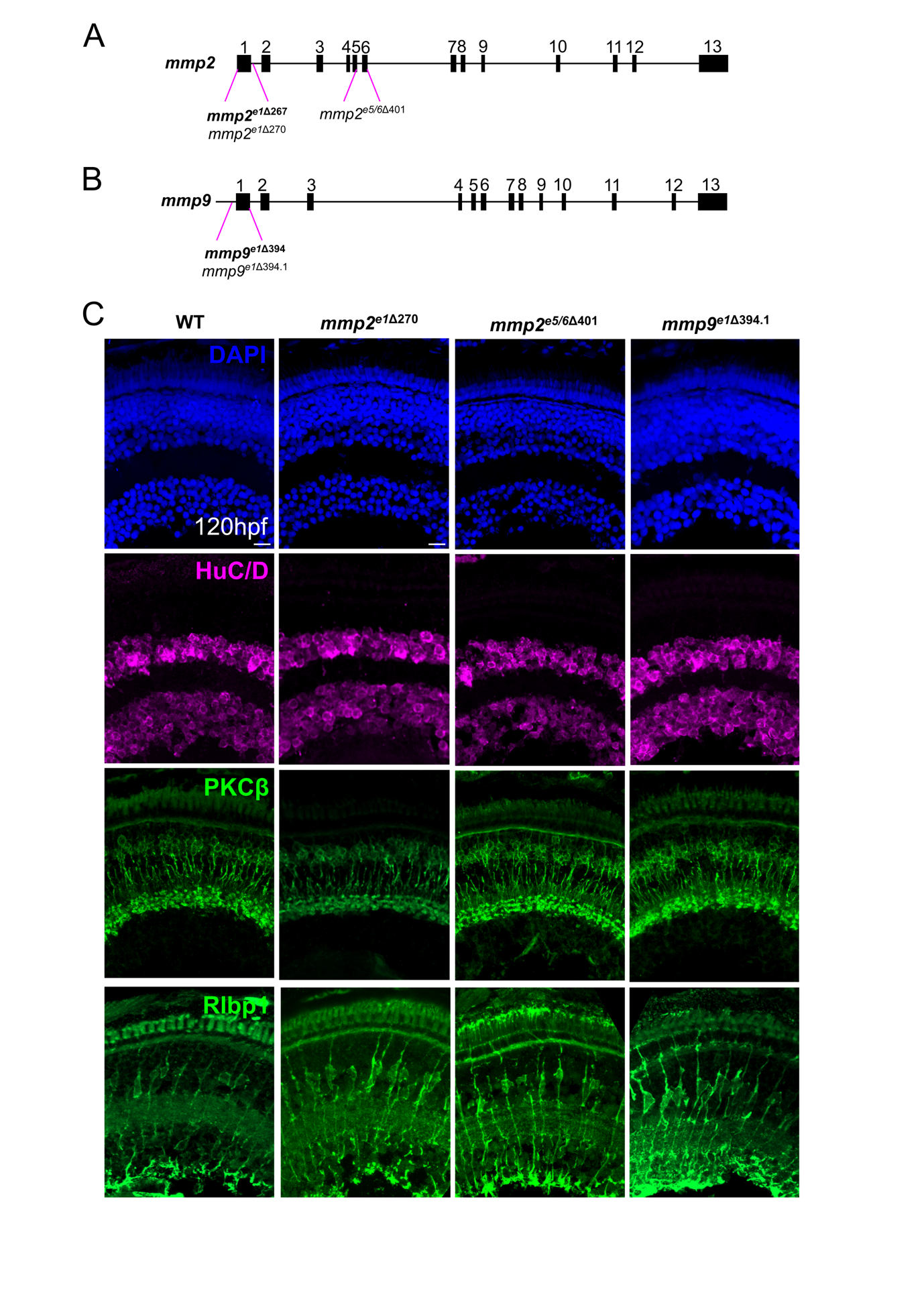
**FIGURE S4**

**Figure S4. Summary of the *mmp2* and *mmp9* mutant alleles generated in this study. (A)** Exon structure of the zebrafish *mmp2* gene highlighting the location of the 3 alleles of *mmp2* mutants raised: *mmp2^e1delta267^* , mmp2^e1270^ and mmp2^e5/6401^ in exons 1 and the latter 5/6 encoding the catalytic domain; Allele highlighted in bold was used in the study. **(B)** Exon structure of the zebrafish *mmp9* gene highlighting the location of the 2 alleles of *mmp2* mutants raised, *mmp9^e1delta394^* and *mmp2^e5^*^/6394.1^ in exon1; Allele highlighted in bold was used in the study. **(C)** IF for neuronal markers HuC/D to label ACs and RGCs (magenta), PKCb to label BPCs (green) and Rlbp1 to label Müller glia (green; lower panel) in the additional alleles not used in the main study. Nuclei labelled with DAPI (blue). Scale bars, 10µm.

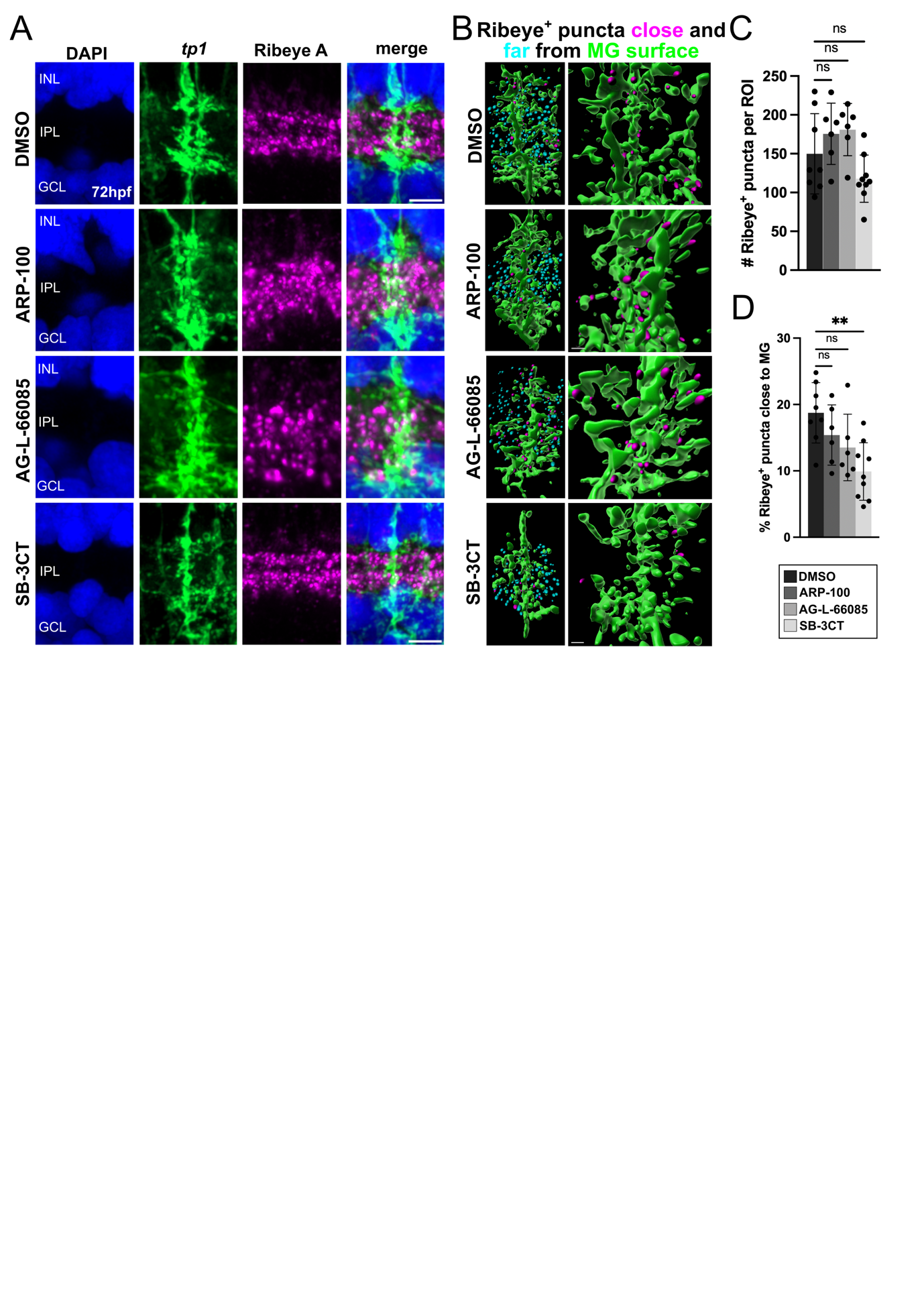

**Figure S5. Formation of Müller glia-synapse interactions is perturbed following pan-gelatinase inhibition. (A)** Close up of the IPL following treatment with DMSO, ARP-100, AG-L-66085 or SB-3CT from 48 to 72 hpf. **(B)** 3D reconstruction of Müller glia IPL process surfaces and Ribeye+ spots categorised as close (magenta) or far (cyan) from Müller glial processes. **(C)** Quantification of the mean number of Ribeye puncta per ROI following treatment with DMOS, ARP-100, AG-L-66085 or SB-3CT. **(D)** Quantification of percentage of Ribeye puncta close to Müller glial processes per ROI following treatment with DMSO, ARP-100, AG-L-66085 or SB-3CT.

**SUPPLEMENTARY TABLES**

**TABLE S1. Transgenic zebrafish lines used in the study.**

| Transgenic line | Promoter specificity in the retina | Authors |
| --- | --- | --- |
| Tg(*tp1:*Venus-Pest)^s940^ | Progenitors, Müller glia | Ninov, Borius and Stainier, 2012 ^105^ |
| Tg(*tp1:*eGFP-CAAX) | Progenitors, Müller glia membranes | Kugler et al. 2023^53^ |
| Tg(*tp1*:H2b-mCherry)^s939^ | Progenitors, Müller glia nuclei | Ninov, Borius and Stainier, 2012 ^105^ |
| Tg(*hsp70:*EMMA-Mmp2) | Ubiquitous, heat-shock inducible | Jeffrey and Crawford, 2018 ^56^ |
| Tg(*ptf1a*:dsRed)^ia6^ | Amacrine and horizontal cells | Yusuf et al. 2012 ^106^ |
| Tg(*vsx1:*GFP)^nns5^ | Bipolar cells | Kimura, Satou and Higashima, 2008 ^107^ |

**TABLE S2. List of antibodies and dilutions used for immunofluorescence**

| Antibody | | | Manufacturer/Supplier | Catalogue # | Host species | | Dilution on cryosections | | | | Dilution in wholemount |
| --- | --- | --- | --- | --- | --- | --- | --- | --- | --- | --- | --- |
|  | anti-Glutamine Synthetase (GS) | | ProteinTech | 66323-1-Ig | Mouse | 1:500 | | | | 1:500 | |
|  | anti-Retinaldehyde binding protein (Rlbp1) | | ProteinTech | 17939-1-AP | Rabbit | 1:300 | | | | 1:200 | |
|  | anti-HuC/D | | Invitrogen | A21271 | Mouse | 1:200 | | | 1:200 | | |
|  | anti-Protein Kinase C-β (PKCβ) | | Proteintech | 12919-1-AP | Rabbit | 1:200 | | | 1:200 | | |
|  | anti-Collagen18-α1 (Col18α1) | | Proteintech | 18301-1-AP | Rabbit | 1:200 | | | 1:100 | | |
|  | anti-Integrin-β1 (Itgβ1) | | Proteintech | 66315-1-Ig | Mouse | 1:200 | | | 1:100 | | |
|  | | anti-Decorin (Dcn) | Proteintech | 14667-1-AP | Rabbit | | | 1:100 | | | n/a |
|  | | anti-Laminin-α1 (Lama1) | Sigma | L9393-2ML | Rabbit | | | 1:100 | | | 1:100 |
|  | | anti-Matrix metalloproteinase 2 (Mmp2) | Santa Cruz | Sc-13594 | Mouse | | | 1:100 | | | 1:100 |
|  | | anti-Matrix metalloproteinase 9 (Mmp9) | Vertebrate Antibodies | V6P3F11*A4 | Mouse | | | 1:30 | | | 1:20 |
|  | | anti-Tissue inhibitor of metalloproteinases 2 (Timp2) | Vertebrate Antibodies | 3A4 | Mouse | | | 1:30 | | | 1:20 |
|  | | anti-4c4 | Gift from A. McGown | n/a | Mouse | | | 1:200 | | | n/a |
|  | | anti-Lcp1 | GeneTex | GTX124420 | Rabbit | | | 1:200 | | | n/a |
|  | | anti-Green fluorescent protein (GFP) | Invitrogen | A11122 | Chicken | | | 1:500 | | | 1:500 |
|  | | anti-Ribeye A | Gift from Teresa Nicholson (Sheets et al., 2012) | n/a | Rabbit | | | 1:500 | | | 1:500 |
|  | | anti-HA | Sigma Aldrich | 11867423001 | Rat | | | n/a | | | 1:200 |
|  | | anti-Rabbit Alexa Fluor^TM^ 546 | Invitrogen | A-11035 | Goat | | | 1:1000 | | | 1:500 |
|  | | anti-Mouse Alexa Fluor^TM^ 647 | Invitrogen | A-21235 | Goat | | | 1:1000 | | | 1:500 |
|  | | anti-Chicken Alexa Fluor^TM^ 488 | Invitrogen | A-11039 | Goat | | | 1:1000 | | | 1:500 |
|  | | anti-Rabbit Alexa Fluor^TM^ 488 | Invitrogen | A-32731 | Goat | | | 1:1000 | | | 1:500 |
|  | | anti-Rat DyLight^TM^ 550 | Invitrogen | SA5-10019 | Goat | | | n/a | | | 1:500 |
|  | | DAPI | Invitrogen | D1306 | n/a | | | 1:1000 | | | 1:500 |
|  | | Phalloidin- Alexa Fluor^TM^ 647 | Invitrogen | A-30107 | n/a | | | n/a | | | 1:500 |

**TABLE S3. Summary of pharmacological MMP inhibitors used in the study.**

| Pharmacological agent | Manufacturer/  Supplier | Catalogue # | Concentration | Target |
| --- | --- | --- | --- | --- |
| SB-3CT | Merck Life Sciences | S1326-5MG | 10 µM | Mmp2 and Mmp9 inhibitor |
| ARP-100 | Generon | A15292-5 | 2 µM | Mmp2 inhibitor |
| AG-L-66085 | Calbiochem/  Sigma Aldrich | 1177749-58-4 | 4 µM | Mmp9 inhibitor |

**TABLE 4. CRISPR-Cas9 crRNA and genotyping primer information.**

| Gene | Primer/crRNA sequence | Amplicon size |
| --- | --- | --- |
| *mmp2* | F primer) **5’-** TTTCACTGCTTTCCCAAATTCT **-3’**  R primer) **5’-** CGGGAACCAAAATGACAATAAT **-3’**  crRNA_1_) AGTCGAACAAACTTGGAGC**TTG**  crRNA_2_) CACTGCTTACGCCGTTCTAA**CGG** | WT:  567 bp  Mutant :  266 bp |
| *mmp9* | F primer) **5’-** CACACTTAATGTGAGATTGCTGG **-3’**  R primer) **5’-** GCGGAGCTATAACATCCAGTT **-3’**  crRNA_1_) TAGTGGGCAAGTGCTGTTAG**TGG**  crRNA_2_) TTTAAGTGTAATTGTCTTCC**AGG** | WT:  791 bp  Mutant:  397 bp |

**TABLE 5. RT-qPCR primer sequences spanning exon-exon junctions.**

| Gene | Primer sequence |
| --- | --- |
| *actb1* | **5’-**CGAGCTGTCTTCCCATCCA**-3’**  **5’-**TCACCAACGTAGCTGTCTTTCTG**-3’** |
| *mmp2* | **5’-** TTTCACTGCTTTCCCAAATTCT **-3’**  **5’-** AAGCTCACTCCAGAACGTGG **-3’** |
| *mmp9* | **5’-** TTCACACGCCTCTTTGACGG **-3’**  **5’-** CACCTGGAGGATAAGCGTGA **-3’** |
